# Antibody evasion by the Brazilian P.1 strain of SARS-CoV-2

**DOI:** 10.1101/2021.03.12.435194

**Authors:** Wanwisa Dejnirattisai, Daming Zhou, Piyada Supasa, Chang Liu, Alexander J. Mentzer, Helen M. Ginn, Yuguang Zhao, Helen M.E. Duyvesteyn, Aekkachai Tuekprakhon, Rungtiwa Nutalai, Beibei Wang, Guido C. Paesen, César López-Camacho, Jose Slon-Campos, Thomas S. Walter, Donal Skelly, Sue Ann Costa Clemens, Felipe Gomes Naveca, Valdinete Nascimento, Fernanda Nascimento, Cristiano Fernandes da Costa, Paola C. Resende, Alex Pauvolid-Correa, Marilda M. Siqueira, Christina Dold, Robert Levin, Tao Dong, Andrew J. Pollard, Julian C. Knight, Derrick Crook, Teresa Lambe, Elizabeth Clutterbuck, Sagida Bibi, Amy Flaxman, Mustapha Bittaye, Sandra Belij-Rammerstorfer, Sarah Gilbert, Miles W. Carroll, Paul Klenerman, Eleanor Barnes, Susanna J. Dunachie, Neil G. Paterson, Mark A. Williams, David R. Hall, Ruben J. G. Hulswit, Thomas A. Bowden, Elizabeth E. Fry, Juthathip Mongkolsapaya, Jingshan Ren, David I. Stuart, Gavin R. Screaton

## Abstract

Terminating the SARS-CoV-2 pandemic relies upon pan-global vaccination. Current vaccines elicit neutralizing antibody responses to the virus spike derived from early isolates. However, new strains have emerged with multiple mutations: P.1 from Brazil, B.1.351 from South Africa and B.1.1.7 from the UK (12, 10 and 9 changes in the spike respectively). All have mutations in the ACE2 binding site with P.1 and B.1.351 having a virtually identical triplet: E484K, K417N/T and N501Y, which we show confer similar increased affinity for ACE2. We show that, surprisingly, P.1 is significantly less resistant to naturally acquired or vaccine induced antibody responses than B.1.351 suggesting that changes outside the RBD impact neutralisation. Monoclonal antibody 222 neutralises all three variants despite interacting with two of the ACE2 binding site mutations, we explain this through structural analysis and use the 222 light chain to largely restore neutralization potency to a major class of public antibodies.

## Introduction

For more than a year SARS-CoV-2 has caused enormous global dislocation, leading to more than 2.5 million deaths (https://www.worldometers.info/coronavirusAccessed:2021-03-01) and leaving no country untouched. Successive waves of infection have led to the imposition of draconian lock downs in many countries resulting in severe economic and societal disruption (Donthu and Gustafsson, 2020).

Enormous investment has been made in vaccine development with hundreds of vaccine candidates in different stages of development using an array of different platforms from RNA, viral vectors, recombinant protein and inactivated (Krammer, 2020). Five vaccines have now been through large scale phase III trials and have demonstrated safety and efficacy (Polack et al., 2020;Voysey et al., 2020;Baden et al., 2020). Four of these, BNT162b2 (Pfizer-BioNTech; mRNA), mRNA-1273 (Moderna; mRNA), ChAdOx1 nCoV-19 (AZD1222) (Oxford-AstraZeneca; chimpanzee adenoviral vectored) and Ad26.COV2-S (Janssen; adenovirus serotype 26 vectored) have received emergency use authorization (EUA) in a variety of countries and are being rolled out at massive scale and NVX-CoV2373 (Novavax; recombinant protein) has also shown impressive efficacy and is likely to achieve EUA in the near future (https://www.medscape.com/viewarticle/944933Accessed:2021-03-01). All of these vaccines have been designed to raise antibodies (and T-cells) to spike protein (S) and because of the speed of development they all include S sequences derived from the first reported sequence from Wuhan in January 2020 (Lu et al., 2020).

SARS-CoV-2 like all RNA viruses has an error prone RNA polymerase, and despite some error correction, progressive accrual of mutational changes is inevitable. The massive scale of the pandemic which is largely uncontrolled leads to huge levels of viral replication, increasing the chances that adaptive mutations will occur. There are many possible ways whereby a mutation in SARS-CoV-2 may give the virus a selective advantage, however concentrating on mutation in S, there are two clear possibilities: increased efficiency of transmission and escape from neutralizing antibodies (Volz et al., 2021).

The S protein is a large Type-1 transmembrane glycoprotein which assembles into homo-trimers (Walls et al., 2020), which form most of the outer surface of coronaviruses. S is divided into two portions S1 and S2 which are cleaved by proteolysis. S1 is responsible for target cell engagement, whilst S2 completes membrane fusion allowing the viral RNA access to the host cell cytoplasm where viral replication can begin. S1 contains an N terminal domain (NTD) and receptor binding domain (RBD).

The RBD interacts with the cellular receptor angiotensin converting enzyme-2 (ACE2) which is expressed on diverse cell types, including cells in the upper and lower respiratory tracts, allowing SARS-CoV-2 to cause respiratory infection. The ACE2 interaction surface is a small 25 amino acid patch at the apex of spike, presented to ACE2 when the RBD swings upwards (Hoffmann et al., 2020;Shang et al., 2020) and it is mutations in this region that are causing the most concern. Three multiply mutated viral strains appeared independently at the end of 2020 in different regions where they rapidly expanded to become the dominant strains (https://www.cogconsortium.uk/wp-content/uploads/2021/01/Report-2_COG-UK_SARS-CoV-2-Mutations.pdf). It is not clear how these strains acquired so many changes without clear intermediate variants. It has however been speculated, with some evidence, that they may have evolved in immunosuppressed chronically infected patients (Kemp et al., 2021) who support high levels of viral replication for months and may be treated with immune plasma or monoclonal antibodies which may drive selection of variants displaying mutations that evade antibody responses.

Of most concern are changes in the RBD. P.1 has three: K417T, E484K and N501Y, B.1.351 also has three: K417N, E484K and N501Y whereas, B.1.1.7 contains the single N501Y mutation. All of these changes have the potential to modulate ACE2/RBD affinity potentially leading to increased transmissibility, for which there is now good evidence in B.1.1.7. In addition, these mutated residues also have the potential to modulate neutralization of SARS-CoV-2 by naturally or vaccine induced antibody responses.

In this paper we examine an isolate of P.1 cultured from a throat swab taken from an infected patient in Manaus, Brazil in December 2020 and compare its interactions with serum and antibodies with those of three other viruses, an early isolate, B.1.1.7 and B.1.351. We test the ability of immune sera induced by infection with early strains of SARS-CoV-2 (Dejnirattisai et al., 2021), or by vaccination with the Oxford-AstraZenca or Pfizer-BioNTech vaccines to neutralize P.1 (Supasa et al., 2021;Zhou et al., 2021). We see a reduction in the neutralizing capacity of immune serum to P.1 similar to the reduction seen with B.1.1.7, but not as severe as that seen with B.1.351 (Zhou et al., 2021). We demonstrate an increased affinity of P.1 RBD for ACE2 and investigate the structural basis of this through crystallography. We also study neutralization by a panel of potent monoclonal antibodies which block RBD/ACE2 interaction and provide a crystallographic solution of how one potent antibody, mAb 222, of the panel (Dejnirattisai et al., 2021) which contacts both K417 and N501, is resistant to the 501Y and 417T/N mutations found in the P.1/B1.351 strains. We dissect the basis for this via a series of high resolution structures of RDB-Fab complexes and based on this restore neutralization of certain antibodies by swapping the light chain. Finally, we bring together data on P.1, B.1.351 and B.1.1.7 and attempt to interpret the different effects these have upon the neutralizing capacity of serum generated to early SARS-CoV-2 strains.

## Results

### The P.1 lineage

P.1 was first reported in December 2020 from Manaus in Amazonas province of Northern Brazil (Faria et al., 2021). A large first wave of infection was seen in Manaus in March-June 2020 and by October around 75% of individuals from the region are estimated to have been infected, representing a very high attack rate. A second large wave of infection began in December 2020 leading to further hospitalizations. This second wave corresponded with the rapid emergence of P.1, not seen before December when it was found in 52% of cases, rising to 85% by January 2021 (**Figure S1**).

P.1 contains multiple changes compared to B.1.1.28 and P.2 which had been previously circulating in Brazil (Faria et al., 2021). Compared to the Wuhan sequence P.1 contains the following mutations: L18F, T20N, P26S, D138Y, R190S in the NTD, K417T, E484K, N501Y in the RBD, D614G and H655Y at the C-terminus of S1 and T1027I, V1176F in S2. The position of the changes seen in P.1 compared with those found in B.1.1.7 and B.1.351 together with a representation on where they occur on the full spike protein and RBD are shown in **Figure 1**. Mutations K417T, E484K, N501Y in the ACE2 interacting surface are of the greatest concern because of their potential to promote escape from the neutralizing antibody response which predominately targets this region (**Figure 1D)** (Dejnirattisai et al., 2021). We searched the COVID-19 genomics UK (COG-UK) (Tatusov et al., 2000) and the global initiative on sharing avian influenza data (GISAID) (https://www.gisaid.org) databases. A small number of sequences including the K417T mutation, inclusive of the P.1 lineage, have been observed in sequencing from Japan, France, Belgium, Italy, the Netherlands and Colombia (**Figure S1**).

**Figure 1.**
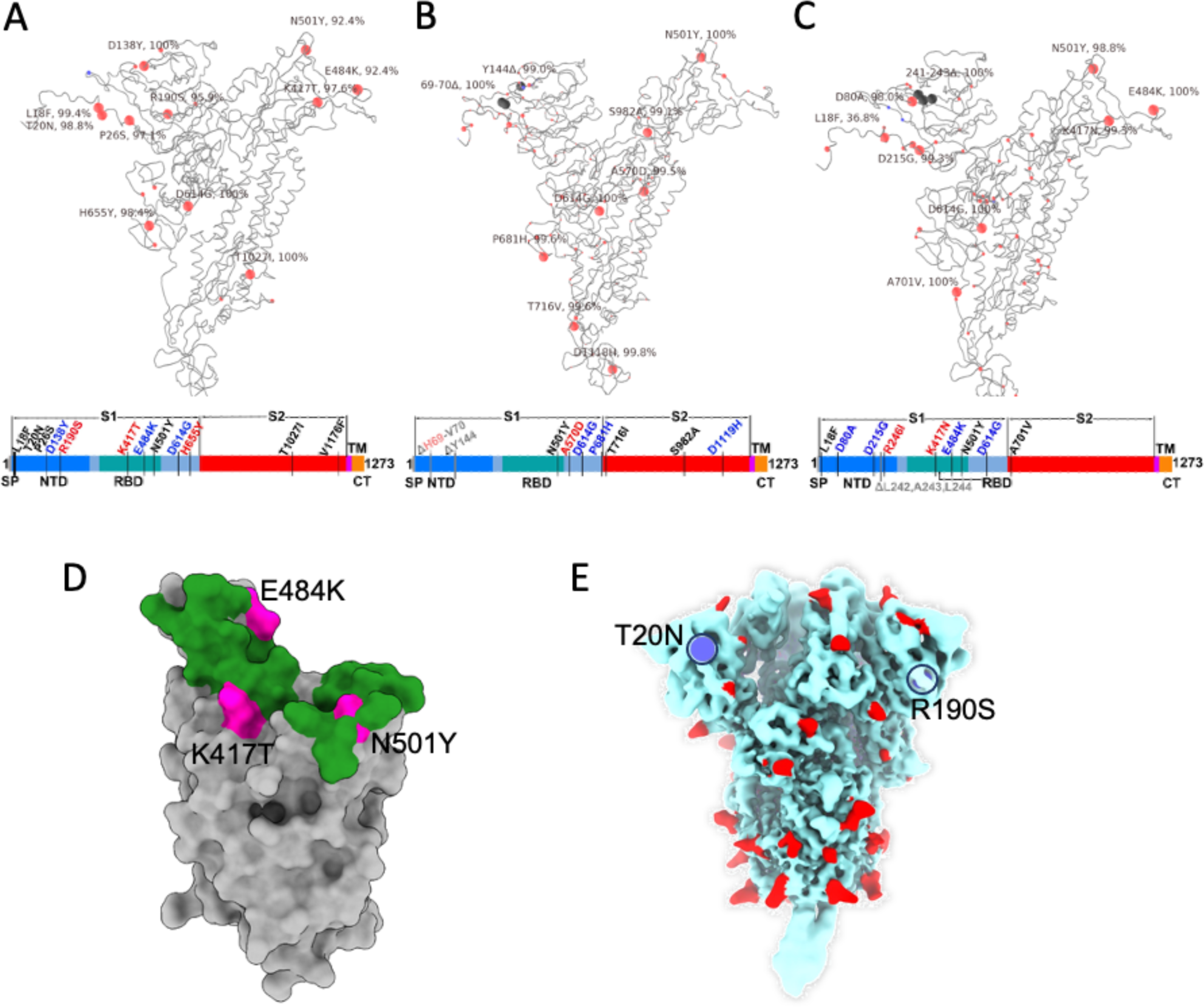
Mutational landscape of P.1. Schematic showing the locations of amino acid substitutions in (A) P.1, (B) B.1.351 and (C) B.1.1.7 relative to the Wuhan SARS-CoV-2 sequence. Under the structural cartoon is a linear representation of S with changes marked on. Where there is a charge change introduced by mutations the change is coloured (red if the change makes the mutant more acidic/less basic, blue more basic/less acidic). (D) Depiction of the RBD as a grey surface with the location of the three mutations K417T, E484K and N501Y (magenta) the ACE2 binding surface of RBD is coloured green. (E) locations of N-linked glycan (red spheres) on the spike trimer shown in a pale blue surface representation, the two new sequons found in P.1 are marked blue.

It is noteworthy that P.1, B.1.1.7 and B.1.351 have accrued multiple mutations in the NTD, in B.1.1.7 there are two deletions *Δ*69-70 and Δ144, in B.1.351 four amino acid changes and the Δ242*−*244 deletion, while in P.1 there are 6 amino acid changes in the NTD but no deletions. Of note, two of the NTD changes in P.1 introduce N-linked glycosylation sequons T20N (TRT to NRT) and R190S (NLR to NLS, **Figure 1E)**. The NTD, in the absence of these changes, reasonably well populated with glycosylation sites, indeed it has been suggested that a single bare patch surrounded by N-linked glycans glycans attached at N17, N74, N122, and N149 defines a ‘supersite’ limiting where neutralizing antibodies can attach to the NTD (Cerutti et al., 2021). Residue 188 is somewhat occluded whereas residue 20 is highly exposed, is close to the site of attachment of neutralizing antibody 159 (Dejnirattisai et al., 2021) and impinges on the proposed NTD supersite.

### The effects of RBD mutations on ACE2 affinity

We have previously measured the affinity of RBD/ACE2 interaction for Wuhan, B.1.1.7 (N501Y) and B.1.351 (K417N, E484K, N501Y) RBDs. N501Y increased affinity 7-fold and the combination of 417, 484 and 501 mutations further increased affinity (19-fold compared to Wuhan). Here we have expressed P.1 RBD (K417T, E484K, N501Y). The K_D_ for the P.1/ACE2 interaction is 4.8 nM with Kon=1.08E5/Ms, Koff=5.18E-4/s (**Figure S2**, Methods), showing that binding to P.1 is essentially indistinguishable from B.1.351 (4.0 nM).

To better understand RBD-ACE2 interactions we determined the crystal structure of the RBD-ACE2 complex at 3.1 Å resolution (Methods, **Table S1**). As expected the mode of RBD-ACE2 engagement is essentially identical for P.1 and the original Wuhan RBD sequence (**Figure 2A**). The RMS deviation between the 791 Ca positions is 0.4 Å, similar to the experimental error in the coordinates, and the local structure around each of the three mutations is conserved. Nevertheless, calculation of the electrostatic potential of the contact surfaces reveals a marked change, with much greater complementarity for the P.1 RBD consistent with higher affinity. (**Figure 2B,C,D**).

**Figure 2.**
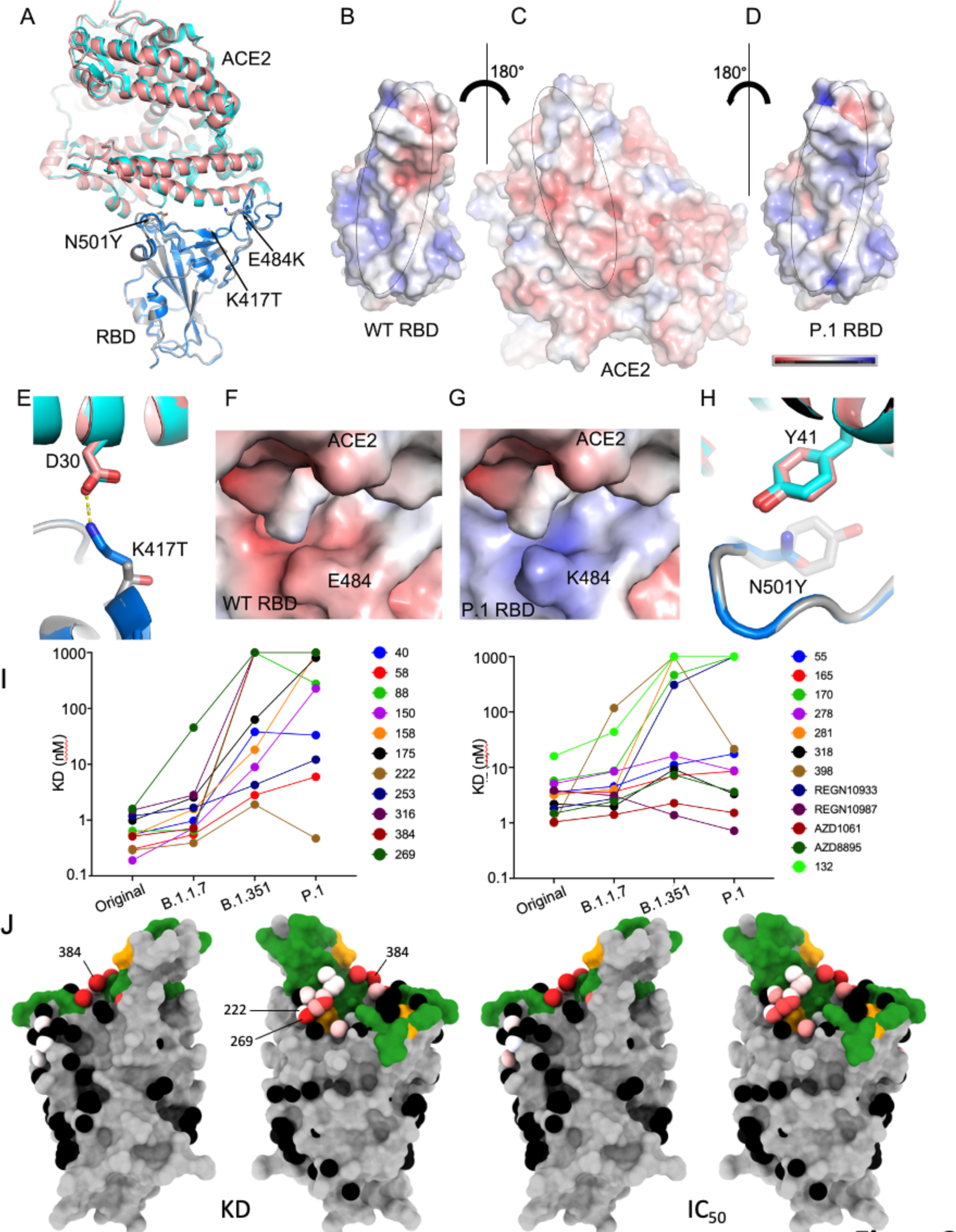
Comparison of WT RBD/ACE2 and P.1 RBD/ACE2 complexes. (A) Comparison of P.1 RBD/ACE2 (grey and salmon) with WT RBD/ACE2 (blue and cyan) (PDB ID 6LZG) by overlapping the RBDs. The mutations in the P.1 RBD are shown as sticks. (B), (C) Open book view of electrostatic surface of the WT RBD/ACE2 complex, and (C), (D) of the P.1 RBD/ACE2 complex. Note the charge difference between the WT and the mutant RBDs. The charge range displayed is ±5 kJ/mol. (E) The K417 of the WT RBD forms a salt bridge with D30 of ACE2. (F) and (G) Effect of E484K mutation on the electrostatic surface (H) Y501 of the P.1 RBD makes a stacking interaction with Y41 of ACE2. (H) K_D_ of RBD/mAb interaction measured by BLI for RBDs of Victoria, B.1.1.7, P.1 and B.1.351 (left to right) (I) BLI data mapped onto the RBD using the method described (Dejnirattisai *et al*., 2021). Front and back views of the RBD are shown. In the left pair the spheres represent the antibody binding sites coloured according to the ratio (K_D_P.1/K_D_Wuhan). For white the ratio is 1, for red it is <0.1 (i.e. at least 10-fold reduction) black dots refer to mapped antibodies not included in this analysis, dark green RBD ACE2 binding surface, yellow mutated K417T, E484K, N501Y. For the right pair atoms are coloured according to the ratio of neutralisation titres (IC50B.1.351/IC50Victoria), for white the ratio is 1, for red it is <0.01 (i.e. at least 100-fold reduction). Note the strong agreement between K_D_ and IC_50_. 269 is very strongly affected and is close to the IGHV3-53 and IGHV3-66 antibodies (*e.g.* 222).

Residue 417 lies at the back of the RBD neck (our RBD anatomy follows Dejnirattisai et al., 2021) and in the original SARS-CoV-2 is a lysine residue which forms a salt-bridge with D30 of ACE2 (**Figure 2E**). The threonine of P.1 RBD no longer forms this interaction and the gap created is open to solvent, so there is no obvious reason why the mutation would increase affinity for ACE2, and this is consistent with directed evolution studies (Zahradník et al., 2021) where this mutation was rarely selected in RBDs with increased affinity for ACE2.

Residue 484 lies atop the left shoulder of the RBD and neither the original Glu nor the Lys of P1 make significant contact with ACE2, nevertheless the marked change in charge substantially improves the electrostatic complementarity (**Figure 2F,G**), consistent with increased affinity.

Residue 501 lies on the right shoulder of the RBD and the change from a relatively short Asn sidechain to the large aromatic Tyr allows for favourable ring stacking interactions consistent with increased affinity (**Figure 2H**).

### Binding of P.1 RBD by potent human monoclonal antibodies

We have previously described a large panel of monoclonal antibodies generated from patients infected with early strains of SARS-CoV-2, before the emergence of B.1.1.7 (Dejnirattisai et al., 2021). From this panel we have selected 20 potent antibodies which have focus reduction neutralization 50% (FRNT50) values <100ng/ml, 19 of these mAbs have an epitope on the RBD and all of these block ACE2/RBD interaction, whilst mAb 159 binds the NTD. We used biolayer interferometry (BLI) to measure the affinity of the RBD-binding antibodies and found that compared to Victoria (SARS-CoV-2/human/AUS/VIC01/2020), an early isolate of SARS-CoV-2, which has a single change S247R in S compared to the Wuhan strain (Seemann et al., 2020; Caly et al., 2020) monoclonal antibody binding was significantly impacted with a number showing complete knock-out of activity (**Figure 2I**). The results with P.1 showed a greater impact compared to B.1.1.7 but similar to B.1.351 (Zhou et al., 2021), this is expected since both contain mutation of the same 3 residues in the RBD, only differing at position 417, K417N in B.1.351 and K417T in P.1. The localization of the impact on binding is shown in **Figure 2J** and reflects direct interaction with mutated residues. Of note is mAb 222 which maintains binding potency across all variants despite adjacency to mutated residues, as discussed below.

### Neutralization of P.1 by potent human monoclonal antibodies

Using the same set of 20 potent antibodies neutralization was measured by a focus reduction neutralization test (FRNT) and compared with neutralization of Victoria and variants B.1.1.7 and B.1.351. Compared to Victoria neutralization by the monoclonal antibodies was significantly impacted by P.1, with 12/20 showing >10-fold reduction in FRNT50 titre and a number showing complete knock out of activity (**Figure 3A Table S2**). The results with P.1 showed a greater impact compared to B.1.1.7 but were as expected similar to those with B.1.351 (Zhou et al., 2021). There is good correlation between the negative impact on neutralization and on RBD-affinity (**Figure 2J**).

**Figure 3.**
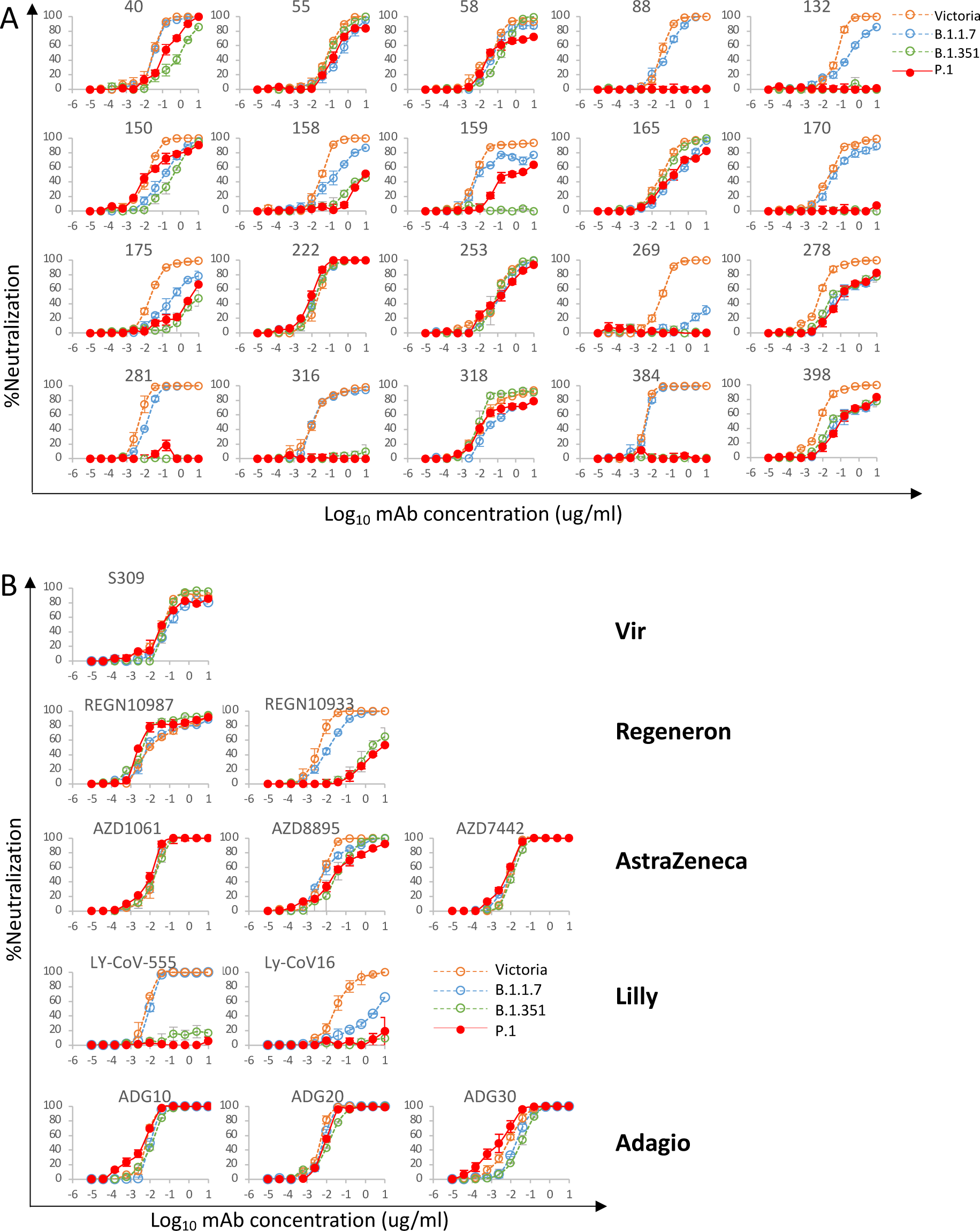
Neutralization of P.1 by monoclonal antibodies. (A) Neutralization of P.1 by a panel of 20 potent human monoclonal antibodies. Neutralization was measured by FRNT, curves for P.1 are superimposed onto curves for Victoria, B.1.1.7 and B.1.351 as previously reported (Supasa et al., 2021;Zhou et al., 2021). FRNT50 titres are reported in Table S2 Neutralization curves for monoclonal antibodies in different stages of development for commercial use. (B) Shows equivalent plots for the Vir, Regeneron, AstraZeneca, Lilly and Adagio antibodies therapeutic antibodies.

### Reduced neutralization of P.1 by monoclonal antibodies being developed for clinical use

A number of potent neutralizing antibodies are being developed for clinical use either therapeutically of prophylactically (Ku et al., 2021;Baum et al., 2020;Kemp et al., 2021). We performed neutralization assays against P.1 using antibodies S309 Vir (Pinto et al., 2020), AZD8895, AZD1061 and AZ7442 (a combination of AZD8895 and AZD1061) AstraZeneca, REGN10987 and REGN10933 Regeneron, LY-CoV555 and LY-CoV16 Lilly and ADG10, ADG20 and ADG30 from Adagio (**Figure 3B**). The affinity of binding to P.1 RBD was also investigated by BLI for the Regeneron and AstraZeneca antibodies and the results (**Figure 2I**) parallel closely the neutralization results. Neutralization of both Lilly antibodies was severely impacted with LY-CoV16 and LY-CoV555 showing almost complete loss of neutralization of P.1 and B.1.351 while LY-CoV16 also showed marked reduction in neutralization of B.1.1.7.

There was also escape from neutralization of P.1 by REGN10933 and a modest reduction in neutralization of P.1 by AZD8895, while AZD1061 and AZD 7442 showed equal neutralization of all SARS-CoV-2 variants. The three Adagio antibodies neutralized all variants with all reaching a plateau at 100% neutralization and interestingly, ADG30 showed a slight increase of neutralization of P.1. S309 Vir was largely unaffected although for several viruses, including P.1, the antibody failed to completely neutralize, conceivably reflecting incomplete glycosylation at N343, since the sugar interaction is key to binding of this antibody N343 (Pinto et al., 2020). The escape from REGN10933 and LY-CoV555 mirrors that of other potent antibodies (including 316 and 384 in our set) which make strong interactions with residues 484-486 and are severely compromised by the marked change E484K, whereas LY-CoV016, an IGHV3-53 mAb, is affected by changes at 417 and 501. The abrogation of the Lilly Ly-CoV-16 and LyCoV-555 antibodies reflects the observation of Starr et al. (Starr et al., 2021)(Greaney et al., 2021) that LY-CoV555 is sensitive to mutation at residue 384 and LY-CoV16 is sensitive to changes at 417.

### Reduced neutralization by an NTD-binding antibody

The neutralization titre of NTD-binding mAb159, was 133-fold reduced on P.1 compared to Victoria with only 64% neutralization at 10μg/ml (**Figures 3A**). Although P.1 does not harbor deletions in the NTD like B.1.1.7 (Δ69*−*70, Δ144) or B.1.351 (Δ242*−*244), it is clear that the 6 NTD mutations in P.1 (L18F, T20N, P26S, D138Y, R190S) disrupt the epitope for mAb159 (**Figure 4A**) (Dejnirattisai et al., 2021;Supasa et al., 2021). It is possible that the failure of this antibody to achieve complete neutralization could be due to partial glycosylation at residue 20, which is some 16 Å from bound Fab 159, however the L18F mutation is even closer and likely to diminish affinity (**Figure 4A**). Since it has been proposed that there is a single supersite for potent NTD binding antibodies we would expect the binding of many of these to be affected (Cerutti et al., 2021).

**Figure 4.**
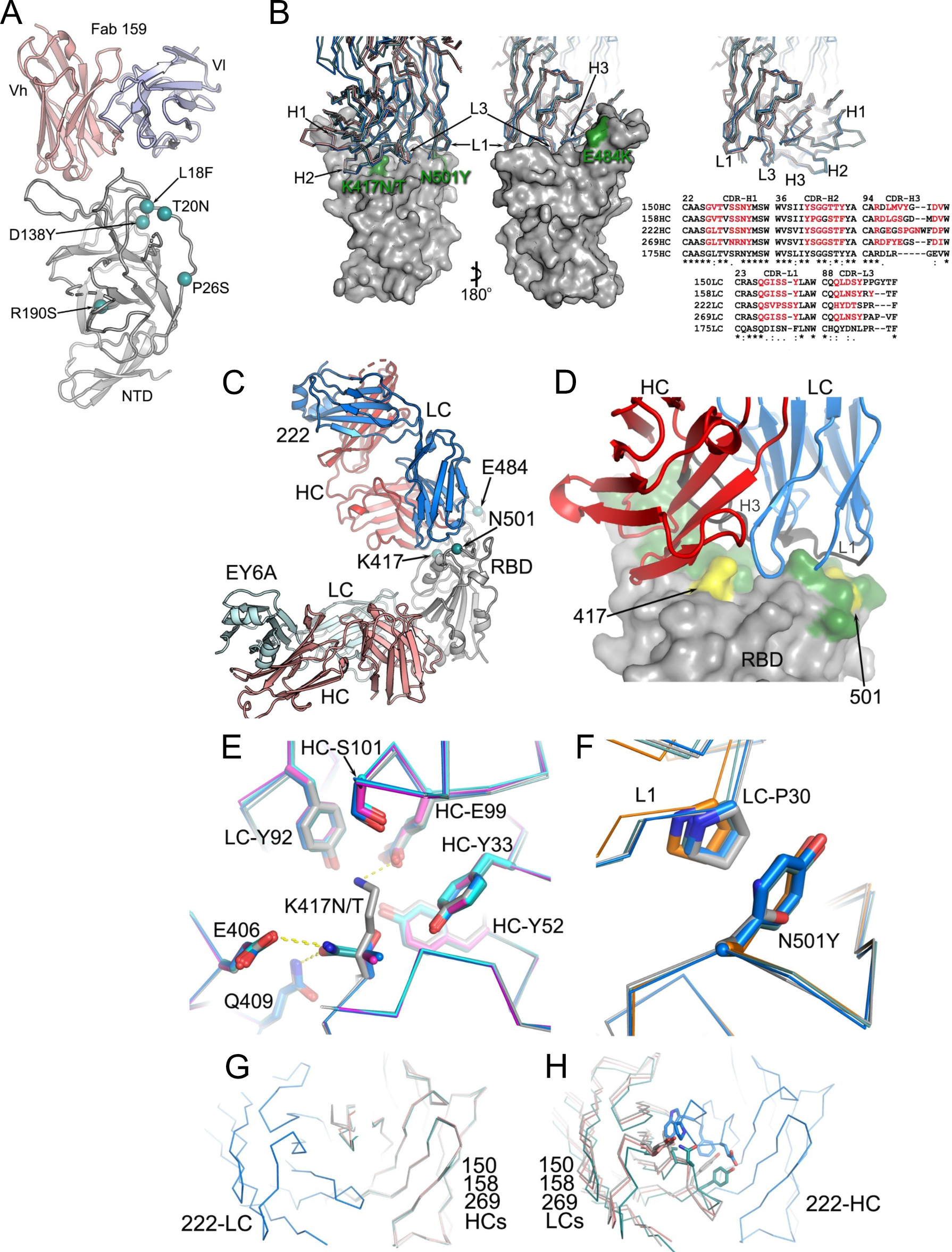
Structures of Fab 222 in complex with P.1 RBD. (A) Ribbon depiction of Fab 159/NTD complex with P1 mutations in the NTD highlighted as cyan spheres. (B) Front and back surfaces of the RBD bound to a typical VH3-53. P1 mutations in the RBD are highlighted in dark green and labelled. In this group, monoclonal antibody 222 has a slightly longer CDR3. Sequences of VH3-53 CDR1-3 heavy and light chains are also shown. (C) Crystal structure of P1 RBD, 222 Fab and EY6A Fab (Zhou et al., 2020). (D) Close up of 222 CDRs interacting with the RBD (grey) mutations are highlighted in yellow on the green ACE2 interface. (E) K417 interactions with Fab 222 (F) N501 interactions with Fab 222. (G), (H) Fab 222 chimera models.

### Reduced neutralization by VH3-53 public antibodies

Five of the potent monoclonal antibodies used in this study (150, 158, 175, 222 and 269), belong to the VH3-53 family and a further 2 (out of 5 of this family) belong to the almost identical VH3-66, and the following discussion applies also to these antibodies. The binding sites for these have been described (Dejnirattisai et al., 2021). The large majority of these antibodies attach to the RBD in a very similar fashion. These motifs recur widely, VH3-53 are the most prevalent deposited sequences and structures for SARS-CoV-2 neutralizing antibodies. Their engagement with the RBD is dictated by CDR-H1 and CDR-H2 whilst the CDR-H3 is characteristically short and makes rather few interactions (Yuan et al., 2020; Barnes et al., 2020;Dejnirattisai et al., 2021). We have previously solved the structures of mAbs 150, 158 and 269 (**Figure 4B**) which show that whilst there are no contacts with residue 484, there are interactions of CDR-H3 with K417 and CDR-L1 with N501, meaning that binding and neutralization by VH3-53 antibodies would be predicted to be compromised by the N501Y change in variant viruses B.1.1.7, B.1.351 and P.1, whilst the additional change at 417 in P.1 (K417T) and B.1.351 (K417N) might be expected to have an additive effect (Dejnirattisai et al., 2021).

In practice, changes in the light chain and CDR-H3 between members of this family mean that the story is more complex. Thus, neutralization of P.1 by 175 and 158 is severely impacted and neutralization of P.1 by 269 is almost completely lost. However, for 150 P.1 neutralization is less compromised than for B.1.351 (Zhou et al., 2021), whilst for 222 neutralization is completely unaffected by the changes in P.1 and indeed all variants (**Figure 3A**).

We measured the affinity of 222 for both P.1 (K_D_ = 1.92±0.01 nM) and Wuhan RBD (K_D_ = 1.36±0.08 nM) showing no reduction in the strength of interaction despite the changes occurring in the putative binding site for P.1 (**Table S2**).

To understand how 222 is able to still neutralize P.1 we solved the crystal structures of six ternary complexes of 222 in complex with the RBDs for (i) the original virus, and bearing mutations (ii) K417N; (iii) K417T; (iv) N501Y; the 417, 484 and 501 changes characteristic of B.1.351 (v) and P.1 (vi). All crystals also contained a further Fab, EY6A as a crystallization chaperone (Zhou et al., 2020), were isomorphous and the resolution of the structures ranged from 1.95 to 2.67 Å, **Figure 4C,D**, Methods, **Table S1**. As expected, the structures are highly similar with the binding pose of 222 being essentially identical in all structures (pairwise RMSD in C*α* atoms between pairs of structures are ∼0.2-0.3 Å for all residues in the RBD and Fv region of mAb 222, **Figure 4D**).

In the original virus residue 417 makes a weak salt bridge interaction with heavy chain CDR3 residue E99. Mutation to either Asn or Thr abolishes this and there is little direct interaction, although there are weak (∼3.5 Å) contacts to heavy chain Y52 and light chain Y92 (**Figure 4E**). However, a buffer molecule/ion rearranges to form bridging interactions and this may mitigate the loss of the salt bridge, in addition the original salt bridge is weak and its contribution to binding may be offset by the loss of entropy in the lysine sidechain. We note that CDR-H3 of 222, at 13 residues is slightly longer than found in the majority of potent VH3-53 antibodies, however this seems unlikely to be responsible for the resilience of 222, rather it seems that there is little binding energy in general from the CDR3-H3, since most of the binding energy contribution of the heavy chain comes from CDR-H1 and CDR-H2 which do not interact with RBD residue 417, meaning that many VH3-53 antibodies are likely to be resilient to the common N/T mutations (**Figure 4B**).

Residue 501 makes contact with CDR-L1 of mAb 222 (**Figure 4D,F**), however the interaction, with P30 is probably slightly strengthened by the N501Y mutation which provides a stacking interaction with the proline, conferring resilience. This is in contrast to the situation with most other VH3.53 antibodies where direct contacts confer susceptibility to escape by mutation to Tyr (**Figures 2I,J and 3A**) (Supasa et al., 2021;Zhou et al., 2021).

### The 222 light chain can rescue neutralization by other VH3-53 mAbs

Reasoning that the relative robustness of mAb 222 to common variants (P.1, B.1.1.7 and B.1.351) compared to other VH3-53 antibodies stems from the choice of light chain we modelled the 222LC with the heavy chains of other VH3-53 antibodies to see if they might be compatible **(Figure 4G)**. The result was striking, it appeared that there would likely be no serious steric clashes. This contrasted with the numerous clashes seen when we docked the light chains of other VH3-53 antibodies onto the heavy chain of 222 (**Figure 4G,H**). This suggests that the 222 light chain might be an almost universal light chain for these 3-53 antibodies and could confer resilience to P.1, B.1.1.7 and B.1.351 variants. This led us to create chimeric antibodies containing the 222LC combined with the HC of the other VH3-53 mAbs 150, 158, 175 and 269. In all cases, chimeric antibodies expressed well and we performed neutralization assays against Victoria, B.1.1.7, B.1.351 and P.1 viruses **(Figure 5)**. For B.1.1.7 neutralization of 150HC/222LC, 158HC/222LC and 269HC/222LC was restored to near the level seen on Victoria, whilst 175HC/222LC could not fully neutralize B.1.1.7. For B.1.351 and P.1 the activity of mAbs 150 and 158 was restored in chimeras containing the 222LC, with the 150HC/222LC showing 50-fold greater potency against B.1.351 (7ng vs 350 ng/ml) and 13-fold greater potency against P.1 (3ng vs 40 ng/ml) than native 150. With an FRNT50 of 3ng/ml 150HC/222LC was the most potent antibody tested against P.1.

**Figure 5.**
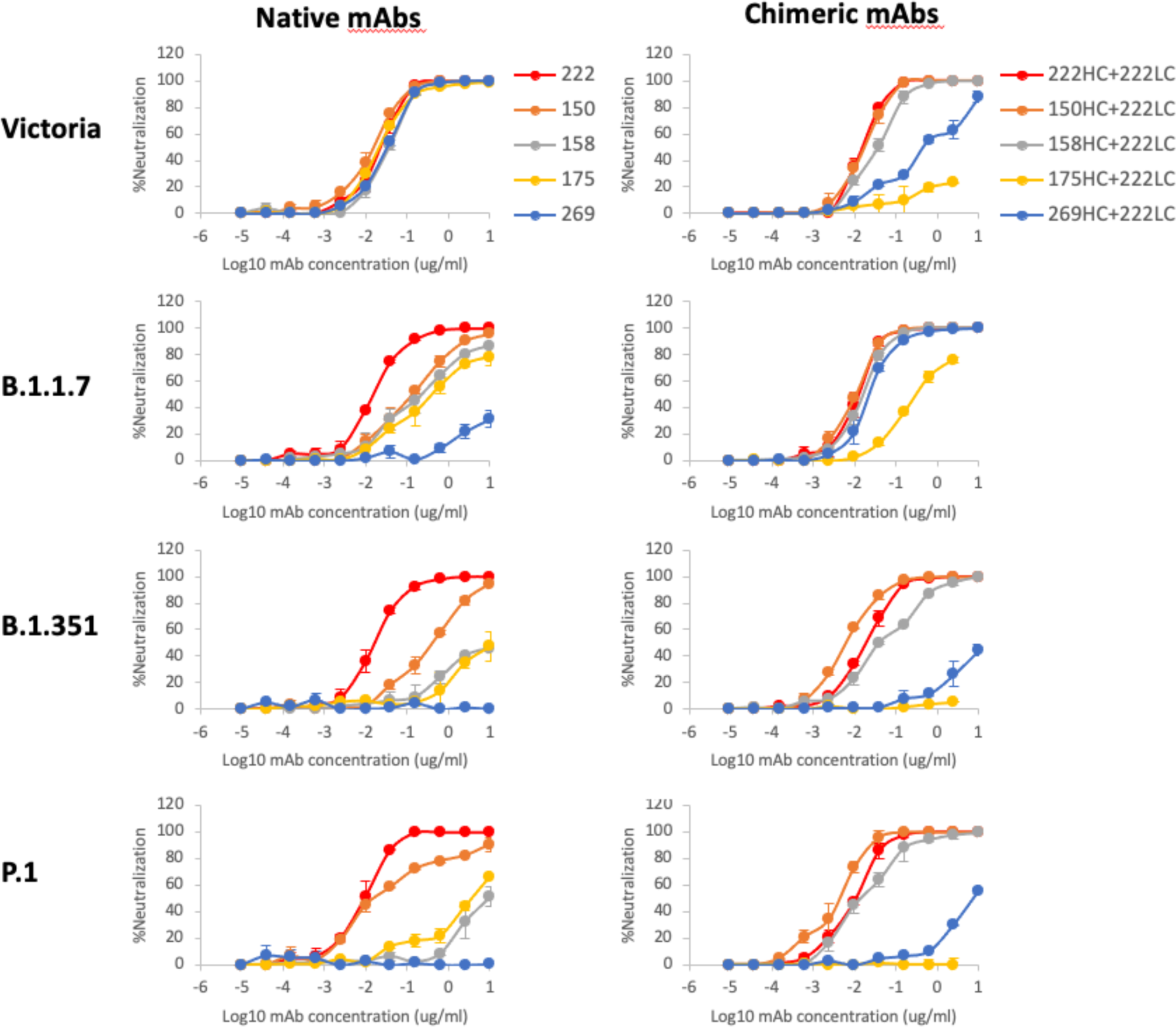
Neutralization curves of VH3-53 chimeric antibodies. Neutralization curves of Victoria, B.1.1.7, B.1.351 and P.1. Left hand column; neutralization curves using the native antibodies 222, 150, 158, 175 and 269. Right hand column; neutralization curves for chimeric antibodies, the heavy chains of 150, 158, 175 and 269 are combined with the light chain of 222, native 222 is used as the control. FRNT50 titres are given in Table S2.

### Neutralization of P.1 by convalescent plasma

We collected convalescent plasma samples from a cohort of volunteers who had suffered from SARS-CoV-2 infection evidenced by a positive diagnostic PCR. Samples were collected during the convalescent phase, 4-9 weeks following infection, all samples were taken during the first wave of infection in the UK, prior June 2020 and well before the emergence of the B.1.1.7 variant. We have also collected plasma from volunteers recently infected with B.1.1.7 as demonstrated by viral sequencing or S gene drop out from the diagnostic PCR (Dejnirattisai et al., 2021;Supasa et al., 2021).

Neutralization of P.1 was assessed by FRNT on 34 convalescent samples **(Figure 6A Table S3A)**. P.1 neutralization curves are displayed alongside neutralization curves for Victoria, together with B.1.1.7 and B.1.351. P.1 geometric mean neutralization titres were reduced 3.1-fold compared to Victoria (p< 0.0001). This reduction was similar to B.1.1.7 (2.9-fold) and considerably less than B.1.351 (13.3-fold) **(Figure 6C)**. When using plasma from individuals infected with B.1.1.7 we saw only modest (1.8-fold p=0.0039) reductions in neutralization comparing P.1 with Victoria **(Figure 6B and D Table S3B)**.

**Figure 6.**
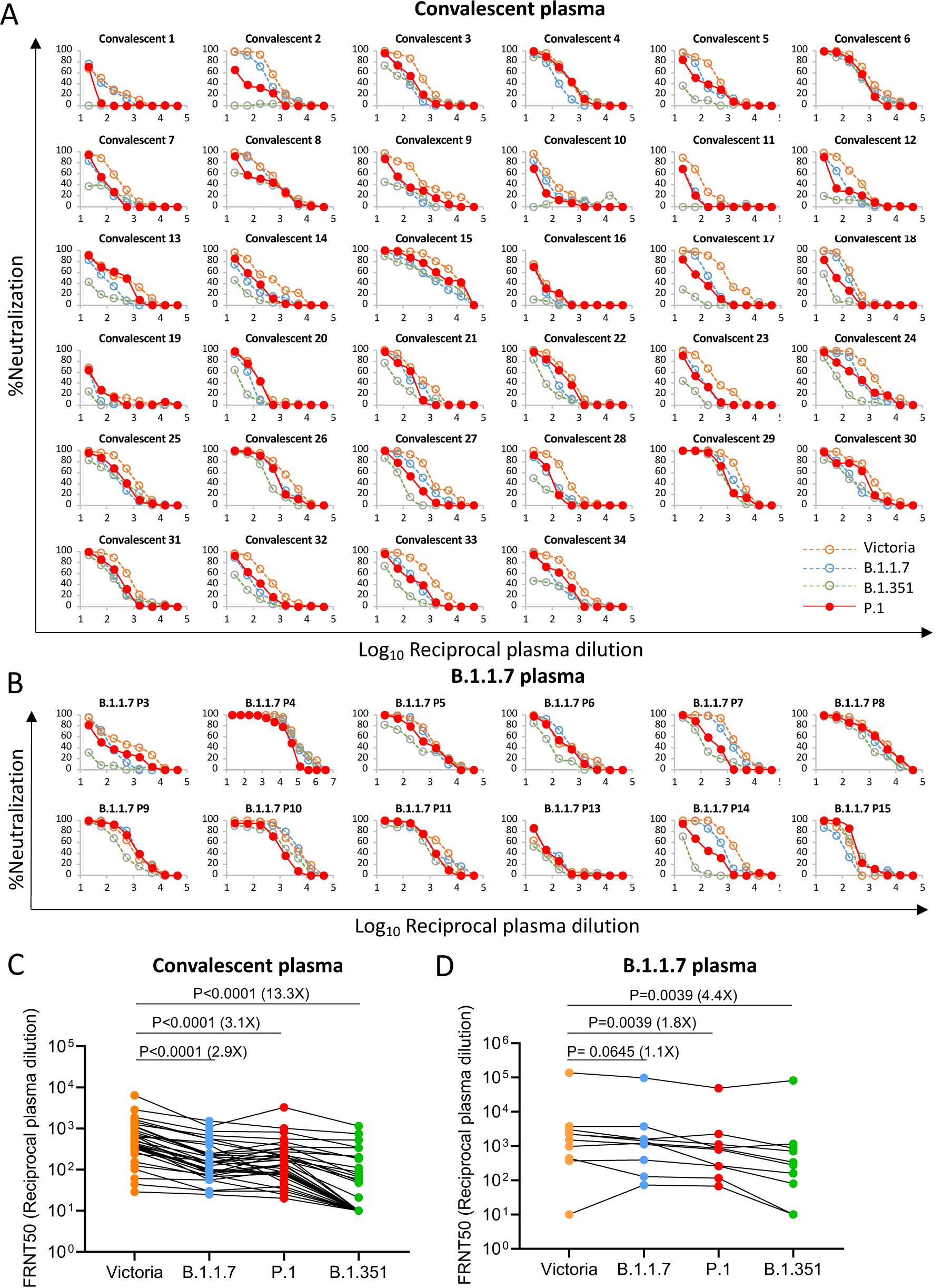
Neutralization of P.1 by convalescent plasma. Plasma (n=34) was collected from volunteers 4-9 weeks following SARS-CoV-2 infection, all samples were collected before June 2020 and therefore represent infection before the emergence of B.1.1.7 in the UK. (A) Neutralization of P.1 was measured by FRNT, comparison is made with neutralization curves for Victoria, B.1.1.7 and B.1.351 that we have previously generated (Zhou et al., 2021;Supasa et al., 2021). (B) Neutralization of P.1 by plasma taken from volunteers who had suffered infection with B.1.1.7 as evidenced by sequencing or S-gene drop out by diagnostic PCR. Samples were taken at varying times following infection. (C-D) Comparison of FRNT50 titres between Victoria and P.1, data for B.1.1.7 and B.1.351 are included for comparison and, the Wilcoxon matched-pairs signed rank test was used for the analysis and two-tailed P values were calculated, geometric mean values are indicated above each column.

### Neutralization of P.1 by vaccine serum

We next performed neutralization assays using serum collected from individuals who had received either the BNT162b2 Pfizer-BioNTech or ChAdOx1 nCoV-19 Oxford-AstraZeneca vaccine **Figure 7** (Supasa et al., 2021;Zhou et al., 2021). For the Pfizer BioNTech vaccine serum was collected 4-14 days following the second dose of vaccine administered three weeks after the first dose (n=25). For the Oxford-AstraZeneca vaccine serum was taken 14 or 28 days following the second dose which was administered 8-14 weeks following the first dose (N=25). Geometric mean neutralization titres against P.1 were reduced 2.6-fold (p<0.0001) relative to the Victoria virus for the Pfizer-BioNTech vaccine serum **Figure 7A,C** and 2.9-fold (P<0.0001) for the Oxford-AstraZeneca vaccine **Figure 7B,D Table S4**.

**Figure 7.**
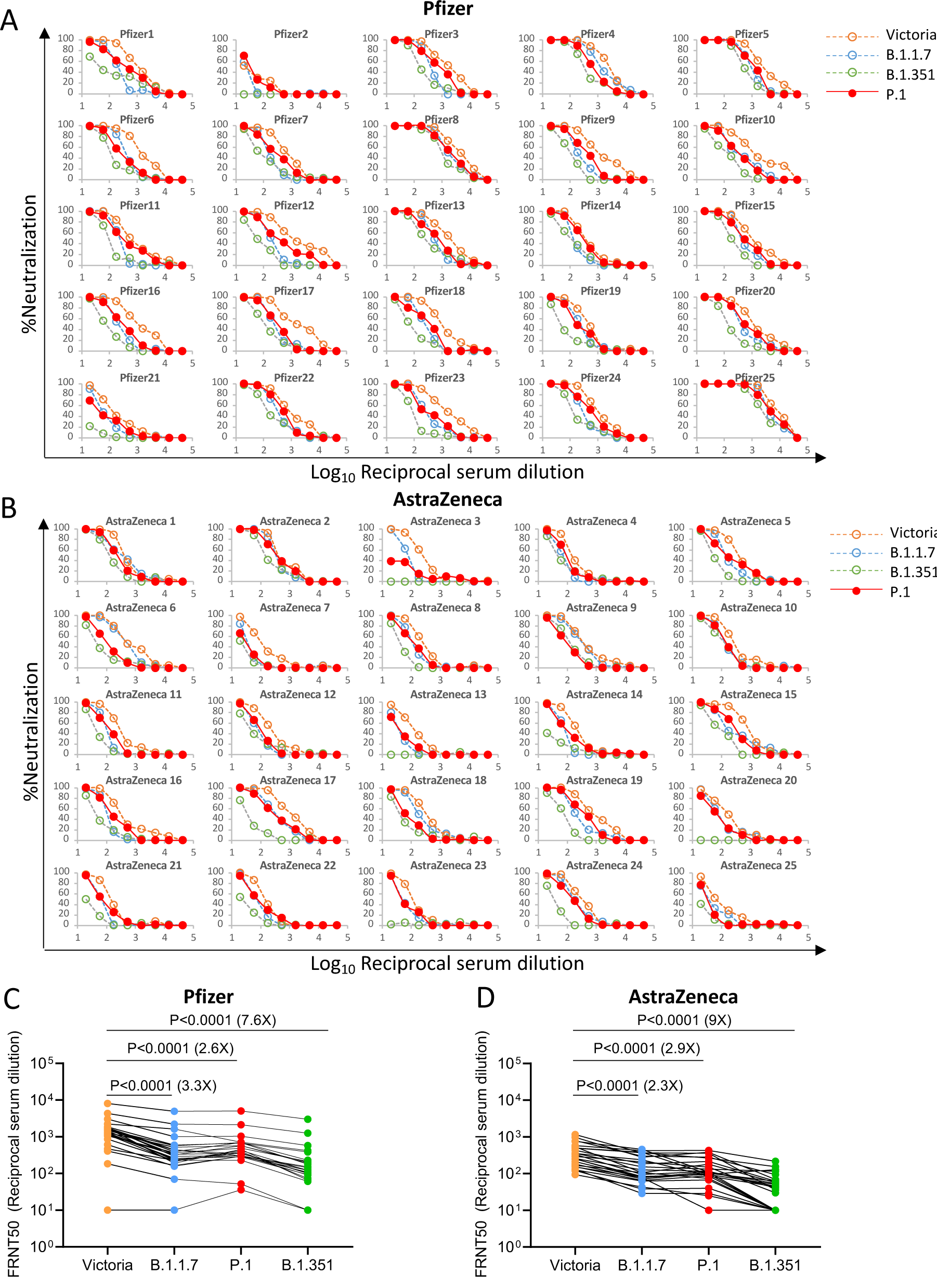
Neutralization of P.1 by vaccine serum. (A) Pfizer vaccine, serum (n=25) was taken 7-17 days following the second dose of the Pfizer-BioNTech vaccine. FRNT titration curves are shown with Victoria, B.1.1.7 and B.1.351 as comparison (Supasa et al., 2021;Zhou et al., 2021). (B) AstraZenca vaccine, serum was taken 14 or 28 days following the second dose of the Oxford-AstraZeneca vaccine (n=25). (C-D) Comparison of FRNT50 titres for individual samples for the Pfizer and AstraZeneca vaccine between Victoria, B.1.1.7, B.1.351 and P.1, the Wilcoxon matched-pairs signed rank test was used for the analysis and two-tailed P values were calculated, geometric mean values are indicated above each column.

Neutralization titres against P.1 were similar to those against B.1.1.7 and only a minority of samples failed to reach 100% neutralization at 1:20 dilution of serum, considerably better than neutralization of B.1.351, where titres were reduced 7.6-fold and 9-fold for the BNT162b2 Pfizer and ChAdOx1 nCoV-19 AstraZeneca vaccines respectively.

## Discussion

Large scale viral sequencing programmes have uncovered a spectrum of mutations containing changes at many locations in the SARS-CoV-2 genome in correspondence with the concept of viral quasispecies (Domingo and Perales, 2019). Mutations in S are of particular concern as S, through the RBD, directs cellular tropism and in addition is the target for the neutralizing antibody response. Mutations in S could therefore enhance viral fitness by increasing affinity to ACE2 or provide escape from the antibody response induced by natural infection or vaccination.

P.1 contains 12 individual changes spread throughout S with three changes in the RBD. In this paper we demonstrate an increase in affinity of interaction for P.1 RBD with ACE2 to an equivalent degree as that observed for B.1.351 with binding somewhat tighter than for B.1.1.7. It seems conceivable that this increase in receptor affinity may drive increased virus transmissibility, allowing the three variants to become dominant strains in the regions where they emerged (Zhou et al., 2021;Supasa et al., 2021).

The ACE2 interacting surface of RBD is a small 25 amino acid patch at the apex of S and is under extreme selection pressure as it not only mediates interaction with the cellular receptor but is also the site of binding for a major class of neutralizing antibodies that block the interaction of ACE2 with the RBD (Zost et al., 2020;Kreye et al., 2020;Wu et al., 2020;Yuan et al., 2020). (Dejnirattisai et al., 2021). Recently, two elegant unbiased approaches have been used to assess the influence of mutation on the ACE2/RBD binding affinity or the ability of RBD mutations to evade the polyclonal antibody response. Firstly, a yeast display approach was used to generate RBD mutants with enhanced ACE2 binding. Amongst a number of mutations selected were the very same positions found in the recent variants of concern, namely E484K and N501Y, and less frequently changes at residue 417 were also observed (Zahradník et al., 2021). Multiple rounds of selection led to the emergence of mutant RBDs with 600-fold higher affinity to ACE2, in the low picomolar range. In a second approach, polyclonal anti-SARS-CoV-2 serum was used to select mutant RBD from a yeast display library which showed reduced antibody binding (Greaney et al., 2021). This approach led to the identification of a number of potential antibody escape mutants, amongst them E484 which is likely responsible for a proportion of the escape from antibody neutralization we describe for P.1.

What is driving the emergence of the new strains is difficult to determine, the emergence of B.1.1.7 occurred on the background of relatively low population immunity and may have been primarily driven by increased transmissibility. The emergence of B.1.351 occurred on the background of around 30% seropositivity in South Africa and P.1 on the background of an estimated 75% seropositivity in Manaus Brazil (Faria et al., 2021). It seems possible that selection of P.1 and B.1.351 may have been in part driven by immune escape, however until methods are developed to screen at a population level for the frequency of reinfection, it is not possible to determine this, especially as reinfection may lead to more mild or asymptomatic disease.

Because P.1 and B.1.351 contain very similar changes in the RBD it might be assumed that neutralization of both would be similarly affected. This was indeed the case for neutralization by monoclonal antibodies directed at the RBD, where there was substantial escape from many antibodies in our panel or from antibodies being developed for clinical use. However, neutralization of P.1 was not compromised as severely as neutralization of B.1.351, when using convalescent or vaccine serum induced by earlier SARS-CoV-2 strains(Zhou et al., 2021). Using convalescent serum B.1.351 showed 13-fold reduction in neutralization compared to Victoria whilst P.1 was only reduced 3.1-fold, comparable to the reduction seen with B.1.1.7, which only harbours the single N501Y change in the RBD (Zhou et al., 2021;Supasa et al., 2021). Similarly, neutralization of P.1 by vaccine serum was less impacted than neutralization of B.1.351 meaning that vaccination with Wuhan S will likely provide some protection against P.1. There is now clinical evidence that the ChAdOx1 nCoV-19 Oxford-AstraZeneca and NVX-CoV2373 Novavax vaccines provide protection from B.1.1.7 (Emary et al., 2021;Mahase, 2021). For B.1.351 both the Novavax (https://www.webmd.com/vaccines/covid-19-vaccine/news/20210131/vaccine-not-as-effective-against-south-african-variant) and Janssen vaccine (https://www.reuters.com/article/us-health-coronavirus-vaccines-johnson-j-idUSKBN29Z0F0) saw a marked decrease in efficacy against B.1.351, but still showed >50% protection against moderate and severe disease, whilst in a Phase II trial, ChAdOx1 nCoV-19 efficacy against mild to moderate disease caused by B.1.351 was 10.4% (95% CI: –76.8; 43.8), but efficacy against severe disease could not be assessed in this study (Madhi et al., 2021).

The reason for the differences in neutralization of B.1.351 and P.1 by immune serum are not immediately clear, but presumably reflect the difference in the mutations introduced outside the RBD. In addition to our mAb 159 a number of potent neutralizing mAbs have been reported that map to the NTD (Cerutti et al., 2021), and this domain has multiple mutations in all three major variant strains: B.1.1.7 has two deletions, B.1.351 has a deletion and four substitutions and P.1 has 6 amino acid substitutions, including the creation of two N-linked glycan sequons (**Figure 1 A-C**). Comparison of neutralization of pseudoviruses expressing only the three RBD mutations (K417N E484K N501Y) of B.1.351 with pseudovirus expressing the full suite of mutations in B.1.351 spike show that the non-RBD changes substantially increase escape from neutralization (Wibmer et al., 2021;Dejnirattisai et al., 2021;Wang et al., 2021). The changes in the NTD of the major variants are far less consistent than those found in the RBD, and there are no strong trends in electrostatic properties (**Figure 1A-C**). It therefore remains unclear what the drivers are for these changes, although one or more of immune escape, co-receptor binding, and modulation of RBD dynamics affecting presentation of the receptor binding site are plausible. Nonetheless, it seems likely that these changes are largely responsible for the non-RBD component of neutralization variation between strains.

A number of public antibody responses have been reported for SARS-CoV-2, principal amongst these being VH3-53/VH3-66 and VH1-58 (VH3-30 is also found but the antibodies are not potent neutralizers) (Yuan et al., 2020;Barnes et al., 2020; Dejnirattisai et al., 2021). We have previously shown that mixing heavy and light chains from antibodies within VH1-58 can increase the neutralization titre by 20-fold from the parent antibodies (chimera of 253HC with 55LC or 165LC) (Dejnirattisai et al., 2021). Here we have shown that chimeras created amongst the VH3-53 antibodies using the 222LC are able to confer broad neutralization to antibodies which have reduced neutralization capacity against the viral variants. Furthermore, the chimera of 150HC with 222LC achieved 13 and 3-fold increases in neutralization titre compared to the parental 150 and 222 mAb respectively. Creation of such antibody chimeras amongst other anti-SARS-CoV2 antibodies may similarly lead to the discovery of more antibodies with enhanced activity. This also suggests that highly effective natural responses against all three variants, and common cross-protective responses, will be found.

The recent emergence of a number of variants of concern has led to efforts to design new vaccines which will be able to protect against the viral variants. Exactly which variants or sequences should be selected is difficult to determine in what is likely to be an evolving situation, as vaccine induced herd immunity increases the selection pressure for immune escape. Based on the results reported here the South African B.1.351 is the variant of greatest concern giving the largest reductions in neutralization titres and evidence of complete failure to neutralize in some cases and we believe developing vaccine constructs to B.1.351 to be the greatest priority.

In summary, we demonstrate that P.1 can escape neutralization by a number of monoclonal antibodies including some being developed for prophylactic or therapeutic use, while other antibodies with epitopes away from the mutated RBD residues retain broad neutralization. Thus S309/AZD1061/REGN10987/ADG10/ADG20/ADG30 showed little to no reduction (<4-fold) in neutralization activity across the three variants, consistent with their previously described broadly neutralizing activities across clade I sarbecoviruses.

In contrast to B.1.351, neutralization of P.1 does not show such a substantial reduction by polyclonal serum induced by natural infection or vaccination and there is no evidence of widespread escape. Despite the reduction in neutralization titres it is hoped that immunization with vaccines designed against parent/ancestral strains will provide protection from P.1.

### Limitations of the study

The in vitro FRNT assays we report here do not measure the effect of complement or antibody dependent cell mediated cytotoxicity which may enhance neutralization in vivo. The role that T cells play in immunity to SARS-CoV-2 and in particular protection from severe disease is unknown and worthy of investigation, but recent findings suggest that CD4 and CD8 T cell responses raised to ancestral strains are minimally impacted by the variants (Alison Tarke et al., 2021;Skelly et al., 2021). It will be interesting to determine the directionality of neutralization between the different variant viruses and naturally acquired antibody responses to them. For instance, there is some suggestion in this report that plasma induced by B.1.1.7 is better able to neutralize B.1.351 and P.1. Measuring neutralization of viral variants by B.1.351 and P.1 serum will give a better idea of cross protection against the other strains.

## Acknowledgements

This work was supported by the Chinese Academy of Medical Sciences (CAMS) Innovation Fund for Medical Science (CIFMS), China (grant number: 2018-I2M-2-002) to D.I.S. and G.R.S. H.M.E.D. and J.Ren are supported by the Wellcome Trust (101122/Z/13/Z), Y.Z. by Cancer Research UK (C375/A17721), TAB and RJGH by the UKRI MRC (MR/S007555/1), D.I.S. and E.E.F. by the UKRI MRC (MR/N00065X/1). D.I.S. is a Jenner Investigator. The National Institute for Health Research Biomedical Research Centre Funding Scheme supports G.R.S. We are also grateful for a Fast Grant from Fast Grants, Mercatus Center to support the isolation of human monoclonal antibodies to SARS-CoV-2 and Schmidt Futures for support of this work. G.R.S. is also supported as a Wellcome Trust Senior Investigator (grant 095541/A/11/Z). This is a contribution from the UK Instruct-ERIC Centre. The Wellcome Centre for Human Genetics is supported by the Wellcome Trust (grant 090532/Z/09/Z). Virus used for the neutralisation assays was isolated by Julian Druce, Doherty Centre, Melbourne, Australia. Chanice Knight, Emily Chiplin, Ross Fothergill and Liz Penn contributed to assays. We acknowledge Diamond Light Source for time on Beamline I03 under Proposal lb27009 for COVID-19 Rapid Access. Huge thanks to the teams, especially at the Diamond Light Source and Department of Structural Biology, Oxford University that have enabled work to continue during the pandemic. The computational aspects of this research were supported by the Wellcome Trust Core Award Grant Number 203141/Z/16/Z and the NIHR Oxford BRC. The Oxford Vaccine work was supported by UK Research and Innovation, Coalition for Epidemic Preparedness Innovations, National Institute for Health Research (NIHR), NIHR Oxford Biomedical Research Centre, Thames Valley and South Midland’s NIHR Clinical Research Network. We thank the Oxford Protective T-cell Immunology for COVID-19 (OPTIC) Clinical team for participant sample collection and the Oxford Immunology Network Covid-19 Response T cell Consortium for laboratory support. We acknowledge the rapid sharing of Victoria, B.1.1.7 and B.1.351 which was isolated by scientists within the National Infection Service at PHE Porton Down. We thank The Secretariat of National Surveillance, Ministry of Health Brazil for assistance in obtaining P.1 samples. This work was supported by the UK Department of Health and Social Care as part of the PITCH (Protective Immunity from T cells to Covid-19 in Health workers) Consortium, the UK Coronavirus Immunology Consortium (UK-CIC) and the Huo Family Foundation. EB and PK are NIHR Senior Investigators and PK is funded by WT109965MA and NIH (U19 I082360). DS is an NIHR Academic Clinical Fellow. The views expressed in this article are those of the authors and not necessarily those of the National Health Service (NHS), the Department of Health and Social Care (DHSC), the National Institutes for Health Research (NIHR), the Medical Research Council (MRC) or Public Health, England.

## Author Information

These authors contributed equally: WD, DZ, PS, CL, AJM.

## Contributions

D.Z. performed BLI interaction analyses. D.Z., J.R., N.G.P., M.A.W. and D.R.H. prepared the crystals, enabled and performed X-ray data collection. J.R., E.E.F., H.M.E.D. and D.I.S. analysed the structural results. G.R.S., J.M., P.S., Y.Z., D.Z., G.C.P, B.W., R.N., A.T., J.S-C., C.L-C. and C.L. prepared the Spike constructs, RBDs, ACE2 and antibodies and, W.D. and P.S. performed neutralization assays. D.C. provided materials. H.M.G. wrote MABSCAPE and performed mapping and cluster analysis, including sequence analyses. S.A.C.C., P. G. N., V.N., F. N., C. F. C., P.C.R., A.P-C., M.M.S., A.J.M., E.B., S.J.D., D.S., C.D., R.L., T.D., A.J.P., J.C.K., P.K., M.W.C., T.L., S.B., A.F., M.B., S.B-R., E.C. and S.G. assisted with patient samples and vaccine trials. E.B., M.C., S.J.D., P.K. and D.S. conceived the study of vaccinated healthcare workers and oversaw the OPTIC Healthcare Worker study and sample collection/processing. G.R.S. and D.I.S. conceived the study and wrote the initial manuscript draft with other authors providing editorial comments. R.J.G.H. and T.A.B. contributed to study design. All authors read and approved the manuscript.

## STAR Methods

### RESOURCE AVAILABILITY

#### Lead Contact

Resources, reagents and further information requirement should be forwarded to and will be responded by the Lead Contact, David I Stuart.

#### Materials Availability

Reagents generated in this study are available from the Lead Contact with a completed Materials Transfer Agreement.

#### Data and Code Availability

The coordinates and structure factors of the crystallographic complexes are available from the PDB with accession codes (see Table S1). Mabscape is available from https://github.com/helenginn/mabscape, https://snapcraft.io/mabscape. The data that support the findings of this study are available from the corresponding authors on request.

### EXPERIMENTAL MODEL AND SUBJECT DETAILS

#### Viral stocks

SARS-CoV-2/human/AUS/VIC01/2020 (Caly et al., 2020), SARS-CoV-2/B.1.1.7 and SARS-CoV-2/B.1.351 were provided by Public Health England, P.1 from a throat swab from Brazil were grown in Vero (ATCC CCL-81) cells. Cells were infected with the SARS-CoV-2 virus using an MOI of 0.0001. Virus containing supernatant was harvested at 80% CPE and spun at 3000 rpm at 4 °C before storage at -80 °C. Viral titres were determined by a focus-forming assay on Vero cells. Victoria passage 5, B.1.1.7 passage 2 and B.1.351 passage 4 stocks were sequenced to verify that they contained the expected spike protein sequence and no changes to the furin cleavage sites. The P.1 virus used in these studies contained the following mutations: L18F, T20N, P26S, D138Y, R190S, K417T, E464K, N501Y, D614G, H655Y, T1027I, V1176F. Passage 1 P.1 virus was sequence confirmed and contained no changes to the furin cleavage site.

#### Bacterial Strains and Cell Culture

Vero (ATCC CCL-81) cells were cultured at 37 °C in Dulbecco’s Modified Eagle medium (DMEM) high glucose (Sigma-Aldrich) supplemented with 10% fetal bovine serum (FBS), 2 mM GlutaMAX (Gibco, 35050061) and 100 U/ml of penicillin–streptomycin. Human mAbs were expressed in HEK293T cells cultured in UltraDOMA PF Protein-free Medium (Cat# 12-727F, LONZA) at 37 °C with 5% CO2. *E.coli DH5α* bacteria were used for transformation of plasmids encoding wt and mutated RBD proteins. A single colony was picked and cultured in LB broth with 50 µg mL^-1^ Kanamycin at 37 °C at 200 rpm in a shaker overnight. HEK293T (ATCC CRL-11268) cells were cultured in DMEM high glucose (Sigma-Aldrich) supplemented with 10% FBS, 1% 100X Mem Neaa (Gibco) and 1% 100X L-Glutamine (Gibco) at 37 °C with 5% CO2. To express RBD, RBD K417T, E484K, N501Y, RBD K417N, RBD K417T, RBD E484K and ACE2, HEK293T cells were cultured in DMEM high glucose (Sigma) supplemented with 2% FBS, 1% 100X Mem Neaa and 1% 100X L-Glutamine at 37 °C for transfection.

#### Participants

Participants were recruited through three studies: Sepsis Immunomics [Oxford REC C, reference:19/SC/0296]), ISARIC/WHO Clinical Characterisation Protocol for Severe Emerging Infections [Oxford REC C, reference 13/SC/0149] and the Gastro-intestinal illness in Oxford: COVID sub study [Sheffield REC, reference: 16/YH/0247]. Diagnosis was confirmed through reporting of symptoms consistent with COVID-19 and a test positive for SARS-CoV-2 using reverse transcriptase polymerase chain reaction (RT-PCR) from an upper respiratory tract (nose/throat) swab tested in accredited laboratories. A blood sample was taken following consent at least 14 days after symptom onset. Clinical information including severity of disease (mild, severe or critical infection according to recommendations from the World Health Organisation) and times between symptom onset and sampling and age of participant was captured for all individuals at the time of sampling.

P.1 virus from throat swabs. The International Reference Laboratory for Coronavirus at FIOCRUZ (WHO) as part of the national surveillance for coronavirus had the approval of the FIOCRUZ ethical committee (CEP 4.128.241) to continuously receive and analyze samples of COVID-19 suspected cases for virological surveillance. Clinical samples (throat swabs) containing P.1 were shared with Oxford University, UK under the MTA IOC FIOCRUZ 21-02.

#### Sera from Pfizer vaccinees

Pfizer vaccine serum was obtained 7-17 days following the second dose of the BNT162b2 vaccine. Vaccinees were Health Care Workers, based at Oxford University Hospitals NHS Foundation Trust, not known to have prior infection with SARS-CoV-2 and were enrolled in the OPTIC Study as part of the Oxford Translational Gastrointestinal Unit GI Biobank Study 16/YH/0247 [research ethics committee (REC) at Yorkshire & The Humber – Sheffield]. The study was conducted according to the principles of the Declaration of Helsinki (2008) and the International Conference on Harmonization (ICH) Good Clinical Practice (GCP) guidelines. Written informed consent was obtained for all patients enrolled in the study. Each received two doses of COVID-19 mRNA Vaccine BNT162b2, 30 micrograms, administered intramuscularly after dilution as a series of two doses (0.3 mL each) 18-28 days apart. The mean age of vaccines was 43 years (range 25-63), 11 male and 14 female.

#### AstraZeneca-Oxford vaccine study procedures and sample processing

Full details of the randomized controlled trial of ChAdOx1 nCoV-19 (AZD1222), were previously published (PMID: 33220855/PMID: 32702298). These studies were registered at ISRCTN (15281137 and 89951424) and ClinicalTrials.gov (NCT04324606 and NCT04400838). Written informed consent was obtained from all participants, and the trial is being done in accordance with the principles of the Declaration of Helsinki and Good Clinical Practice. The studies were sponsored by the University of Oxford (Oxford, UK) and approval obtained from a national ethics committee (South Central Berkshire Research Ethics Committee, reference 20/SC/0145 and 20/SC/0179) and a regulatory agency in the United Kingdom (the Medicines and Healthcare Products Regulatory Agency). An independent DSMB reviewed all interim safety reports. A copy of the protocols was included in previous publications (PMID: 33220855/PMID: 32702298).

Data from vaccinated volunteers who received two vaccinations are included in this paper. Vaccine doses were either 5 × 10^10^ viral particles (standard dose; SD/SD cohort n=21) or half dose as their first dose (low dose) and a standard dose as their second dose (LD/SD cohort n=4). The interval between first and second dose was in the range of 8-14 weeks. Blood samples were collected and serum separated on the day of vaccination and on pre-specified days after vaccination e.g. 14 and 28 days after boost.

### METHOD DETAILS

#### Focus Reduction Neutralization Assay (FRNT)

The neutralization potential of Ab was measured using a Focus Reduction Neutralization Test (FRNT), where the reduction in the number of the infected foci is compared to a negative control well without antibody. Briefly, serially diluted Ab or plasma was mixed with SARS-CoV-2 strain Victoria or P.1 and incubated for 1 hr at 37 °C. The mixtures were then transferred to 96-well, cell culture-treated, flat-bottom microplates containing confluent Vero cell monolayers in duplicate and incubated for a further 2 hrs followed by the addition of 1.5% semi-solid carboxymethyl cellulose (CMC) overlay medium to each well to limit virus diffusion. A focus forming assay was then performed by staining Vero cells with human anti-NP mAb (mAb206) followed by peroxidase-conjugated goat anti-human IgG (A0170; Sigma). Finally, the foci (infected cells) approximately 100 per well in the absence of antibodies, were visualized by adding TrueBlue Peroxidase Substrate. Virus-infected cell foci were counted on the classic AID EliSpot reader using **AID ELISpot software.** The percentage of focus reduction was calculated and IC_50_ was determined using the probit program from the SPSS package.

#### Cloning of ACE2 and RBD proteins

The constructs of EY6A Fab, 222 Fab, ACE2, WT RBD, B.1.1.7 and B.1.351 mutant RBD are the same as previously described (Dejnirattisai et al. 2021, Zhou et al., 2021, Supasa et al., 2021). To clone RBD K417T and RBD K417N, primers of RBD K417T (forward primer 5’-GGGCAGACCGGCACGATCGCCGACTAC-3’ and reverse primer 5’-GTAGTCGGCGATCGTGCCGGTCTGCCC) and primers of RBD K417N (forward primer 5’-CAGGGCAGACCGGCAATATCGCCGACTACAATTAC-3’ and reverse primer 5’-GTAATTGTAGTCGGCGATATTGCCGGTCTGCCCTG-3’) were used separately, together with two primers of pNEO vector (Forward primer 5’-CAGCTCCTGGGCAACGTGCT-3’ and reverse primer 5’-CGTAAAAGGAGCAACATAG-3’) to do PCR, with the plasmid of WT RBD as the template. To clone P.1 RBD, the construct of B.1.351 RBD was used as the template and the primers of RBD K417T and of pNEO vector mentioned above were used to do PCR. Amplified DNA fragments were digested with restriction enzymes AgeI and KpnI and then ligated with digested pNEO vector. All constructs were verified by sequencing.

#### Protein production

Protein production was as described in Zhou et al. (Zhou et al., 2020). Briefly, plasmids encoding proteins were transiently expressed in HEK293T (ATCC CRL-11268) cells. The conditioned medium was dialysed and purified with a 5-ml HisTrap nickel column (GE Healthcare) and further polished using a Superdex 75 HiLoad 16/60 gel filtration column (GE Healthcare).

#### Bio-Layer Interferometry

BLI experiments were run on an Octet Red 96e machine (Fortebio). To measure the binding affinity of ACE2 with P.1 RBD and affinities of monoclonal antibodies and ACE2 with native RBD and, RBD K417N, RBD K417T, RBD E484K and RBD K417T E484K N501Y, eachP.1 RBD, each RBD was immobilized onto an AR2G biosensor (Fortebio). Monoclonal antibodies (Dejnirattisai et al., 2021) were used as analytes or serial dilutions of ACE2 were used as analytes. All experiments were run at 30 °C. Data were recorded using software Data Acquisition 11.1 (Fortebio) and Data Analysis HT 11.1 (Fortebio) with a 1:1 fitting model used for the analysis.

#### Antibody production

AstraZeneca and Regeneron antibodies were provided by AstraZeneca, Vir, Lilly and Adagio antibodies were provided by Adagio. For the chimeric antibodies heavy and light chains of the indicated antibodies were transiently transfected into 293Y cells and antibody purified purified from supernatant on protein A.

#### Crystallization

ACE2 was mixed with P.1 RBD in a 1:1 molar ratio to a final concentration of 12.5 mg ml^−1^. EY6A Fab, 222 Fab and WT or mutant RBD were mixed in a 1:1:1 molar ratio to a final concentration of 7.0 mg ml^−1^. All samples were incubated at room temperature for 30 min. Most crystallization experiments was set up with a Cartesian Robot in Crystalquick 96-well X plates (Greiner Bio-One) using the nanoliter sitting-drop vapor-diffusion method, with 100 nl of protein plus 100 nl of reservoir in each drop, as previously described (Water et al., 2003). Crystallization of B.1.1.7 RBD/EY6A/222 complex was set up by hand pipetting, with 500 nl of protein plus 500 nl of reservoir in each drop. Good crystals of EY6A Fab and 222 Fab complexed with WT, K417T, K417N, B.1.1.7, B.1.351 or P.1 RBD were all obtained from Hampton Research PEGRx 2 screen, condition 35, containing 0.15 M Lithium sulfate, 0.1 M Citric acid pH 3.5, 18% w/v PEG 6,000. Crystals of P.1 RBD/ACE2 complex were formed in Hampton Research PEGRx 1 screen, condition 38, containing 0.1 M Imidazole pH 7.0 and 20% w/v Polyethylene glycol 6,000.

#### X-ray data collection, structure determination and refinement

Crystals of ternary complexes of WT and mutant RBD/EY6A and 222 Fabs and the P.1. RBD/ACE2 were mounted in loops and dipped in solution containing 25% glycerol and 75% mother liquor for a second before being frozen in liquid nitrogen prior to data collection. Diffraction data were collected at 100 K at beamline I03 of Diamond Light Source, UK. All data (except some of the WT RBD-EY6A-222 Fab complex images) were collected as part of an automated queue system allowing unattended automated data collection (https://www.diamond.ac.uk/Instruments/Mx/I03/I03-Manual/Unattended-Data-Collections.html). Diffraction images of 0.1° rotation were recorded on an Eiger2 XE 16M detector (exposure time of either 0.004 or 0.006 s per image, beam size 80×20 μm, 100% beam transmission and wavelength of 0.9763 Å). Data were indexed, integrated and scaled with the automated data processing program Xia2-dials (Winter, 2010;Winter *et al*., 2018). A data set of 1080° was collected from 3 positions of a frozen crystal for the WT RBD-EY6A-222 Fab complex. 720° of data was collected for each of the B.1.1.7, P.1 and B.1.351 mutant RBD/EY6A and 222 Fab complexes (each from 2 crystals), and 360° for each of the K417N and K417T RBD with EY6A and 222 Fabs, and ACE2 complexes was collected from a single crystal.

Structures of WT RBD-EY6A-222 and the P.1 RBD-ACE2 complexes were determined by molecular replacement with PHASER (McCoy et al., 2007) using search models of SARS-CoV-2 RBD-EY6A-H4 (PDB ID 6ZCZ) (Zhou et al., 2020) and RBD-158 (PDB ID, 7BEK) (Dejnirattisai et al., 2021) complexes, and a RBD and ACE2 complex (PDB ID, 6LZG (Wang et al., 2020)), respectively. Model rebuilding with COOT (Emsley and Cowtan, 2004) and refinement with PHENIX (Liebschner et al., 2019) were done for all the structures. The ChCl domains of EY6A are flexible and have poor electron density. Data collection and structure refinement statistics are given in **Table S1**. Structural comparisons used SHP (Stuart et al., 1979), residues forming the RBD/Fab interface were identified with PISA (Krissinel and Henrick, 2007) and figures were prepared with PyMOL (The PyMOL Molecular Graphics System, Version 1.2r3pre, Schrödinger, LLC).

#### Quantification and statistical analysis

Statistical analyses are reported in the results and figure legends. Neutralization was measured by FRNT. The percentage of focus reduction was calculated and IC_50_ was determined using the probit program from the SPSS package. The Wilcoxon matched-pairs signed rank test was used for the analysis and two-tailed P values were calculated and geometric mean values. BLI data were analysed using Data Analysis HT 11.1 (Fortebio) with a 1:1 fitting model.

