## Supplementary tables and figures for "Antibody evasion by the Brazilian P.1 strain of SARS-CoV-2"

#### Table S1 Data collection and refinement statistics of RBD complexes

| Structure | RBD-EY6A-222 | K417N-RBD-EY6A-222 | K417T-RBD-EY6A-222 | B.1.1.7-RBD-EY6A-222 | P.1-RBD-EY6A-222 | B.1.351-RBD-EY6A-222 | P.1-RBD-ACE2 |
|---|---|---|---|---|---|---|---|
| **Data collection** | | | | | | | |
| Space group | $P2_12_12_1$ | $P2_12_12_1$ | $P2_12_12_1$ | $P2_12_12_1$ | $P2_12_12_1$ | $P2_12_12_1$ | $P4_12_12$ |
| Cell dimensions | | | | | | | |
| a, b, c (Å) | 54.4, 120.1, 211.7 | 54.8, 122.5, 214.2 | 54.8, 122.7, 213.9 | 54.3, 120.9, 210.2 | 54.7, 122.8, 212.3 | 54.2, 120.4, 211.6 | 103.5, 103.5, 225.9 |
| a, b, g (°) | 90, 90, 90 | 90, 90, 90 | 90, 90, 90 | 90, 90, 90 | 90, 90, 90 | 90, 90, 90 | 90, 90, 90 |
| Resolution (Å) | 120–2.25 (2.29–2.25)[a] | 62–2.24 (2.28–2.24) | 81–1.95 (1.98–1.95) | 79–2.40 (2.44–2.40) | 80–2.67 (2.72–2.67) | 61–2.50 (2.54–2.50) | 70–3.14 (3.27–3.14) |
| $R_{merge}$ | 0.438 (---) | 0.185 (---) | 0.120 (0.990) | 0.257 (---) | 0.365 (---) | 0.401 (---) | 0.500 (---) |
| $R_{pim}$ | 0.073 (0.767) | 0.053 (0.805) | 0.034 (0.972) | 0.051 (0.547) | 0.072 (1.176) | 0.080 (1.086) | 0.100 (1.087) |
| I/s(I) | 4.3 (0.3) | 5.7 (0.3) | 12.0 (0.4) | 7.7 (0.6) | 7.0 (0.4) | 4.6 (0.3) | 5.3 (0.5) |
| $CC_{1/2}$ | 0.991 (0.32) | 0.992 (0.328) | 0.996 (0.354) | 0.995 (0.584) | 0.997 (0.344) | 0.992 (0.360) | 0.992 (0.306) |
| Completeness (%) | 99.8 (95.8) | 100 (99.1) | 100 (99.6) | 100 (98.5) | 100 (97.7) | 99.9 (94.2) | 100 (98.1) |
| Redundancy | 34.8 (17.3) | 13.3 (13.3) | 13.4 (13.2) | 26.6 (25.8) | 26.3 (27.5) | 26.3 (27.1) | 25.7 (27.1) |
| **Refinement** | | | | | | | |
| Resolution (Å) | 106–2.25 | 62–2.24 | 81–1.95 | 79–2.40 | 80–2.67 | 61–2.50 | 70–3.14 |
| No. reflections | 61666/3310 | 65884/3438 | 100792/5255 | 52327/2727 | 39534/2103 | 45980/2393 | 20278/1904 |
| $R_{work}$ / $R_{free}$ | 0.207/0.247 | 0.221/0.251 | 0.223/0.246 | 0.225/0.247 | 0.224/0.255 | 0.211/0.255 | 0.229/0.276 |
| No. atoms | | | | | | | |
| Protein | 8014 | 8013 | 8017 | 8018 | 8034 | 8017 | 6407 |
| | 386 | 423 | 389 | 298 | 182 | 145 | 71 |
| Ligand/ion/water | | | | | | | |
| B factors (Å$^2$) | | | | | | | |
| Protein | 67 | 84 | 78 | 76 | 93 | 78 | 91 |
| | 60 | 67 | 50 | 67 | 84 | 87 | 107 |
| Ligand/ion/water | | | | | | | |
| r.m.s. deviations | | | | | | | |
| Bond lengths (Å) | 0.002 | 0.002 | 0.002 | 0.003 | 0.002 | 0.002 | 0.003 |
| Bond angles (°) | 0.5 | 0.5 | 0.5 | 0.6 | 0.5 | 0.5 | 0.5 |

Values in parentheses are for highest-resolution shell.

### Table S2

| mAb | IC50 (ug/ml) | | | | $FRNT_{50}$ ratio | | | KD (nM) | | | | | Immunoglobulin gene usage | | |
|---|---|---|---|---|---|---|---|---|---|---|---|---|---|---|---|
| | Victoria | B.1.1.7 | B.1.351 | P.1 | B.1.1.7/ Viictoria | B.1.351/ Victoria | P.1/ Victoria | Native RBD | K417T | K417N | E484K | RBD P.1 | IGHV | K/λ | IGLV |
| 40 | 0.026 ± 0.007 | 0.035 ± 0.008 | 0.738 ± 0.311 | 0.153 ± 0.037 | 1.4 | 28.6 | 5.9 | 1.33±0.03 | 38.90±0.89 | 56.7±1.18 | 7.31±0.03 | 38.20±0.95 | 3-66 | K | 1-33 or 1D-33 |
| 55 | 0.095 ± 0.015 | 0.348 ± 0.044 | 0.127 ± 0.014 | 0.306 ± 0.046 | 3.7 | 1.3 | 3.2 | 2.66±0.05 | 4.60±0.12 | 4.21±0.09 | 3.82±0.02 | 11.10±0.37 | 1-58 | K | 3-20 |
| 58 | 0.041 ± 0.003 | 0.116 ± 0.029 | 0.136 ± 0.010 | 0.236 ± 0.075 | 2.8 | 3.3 | 5.7 | 1.46±0.04 | 1.83±0.02 | 2.75±0.03 | 0.97±0.04 | 2.84±0.03 | 3-9 | λ | 3-21 |
| 88 | 0.033 ± 0.001 | 0.058 ± 0.008 | >10 | >10 | 1.8 | >307.6 | >307.6 | 2.90±0.03 | Knocked out | Knocked out | 5.37±0.06 | Knocked out | 4-61 | λ | 1-36 |
| 132 | 0.048 ± 0.000 | 0.337 ± 0.048 | >10 | >10 | 7.0 | >208.4 | >208.4 | 2.93±0.10 | 1846±364 | 2495±1095 | 29.80±0.36 | 1338±452 | 4-34 | λ | 7-46 |
| 150 | 0.012 ± 0.000 | 0.139 ± 0.019 | 0.350 ± 0.010 | 0.040 ± 0.003 | 12.0 | 30.0 | 3.4 | 0.40±0.01 | 1.64±0.02 | 3.35±0.03 | 1.83±0.03 | 8.90±0.13 | 3-53 | K | 1-9 |
| 158 | 0.031 ± 0.004 | 0.254 ± 0.109 | >10 | >10 | 8.3 | >327.5 | >327.5 | 1.89±0.03 | 4.83±0.06 | 9.35±0.11 | 3.03±0.10 | 18.20±0.31 | 3-53 | K | 1-9 |
| 159 | 0.011 ± 0.000 | 0.061 ± 0.020 | >10 | 1.434 ± 0.804 | 5.7 | >928.4 | 133.2 | N/A | N/A | N/A | N/A | N/A | 3-30 | K | 3-20 |
| 165 | 0.034 ± 0.004 | 0.212 ± 0.004 | 0.054 ± 0.013 | 0.241 ± 0.030 | 6.3 | 1.6 | 7.2 | 2.15±0.03 | 2.62±0.06 | 3.29±0.05 | 3.49±0.03 | 7.10±0.15 | 1-58 | K | 3-20 |
| 170 | 0.025 ± 0.004 | 0.105 ± 0.050 | >10 | >10 | 4.2 | >402.2 | >402.2 | 5.23±0.02 | 6.24±0.04 | 6.95±0.05 | Knocked out | 463.4±25.4 | 5-51 | K | 2D-29 |
| 175 | 0.026 ± 0.000 | 0.575 ± 0.280 | >10 | 3.881 ± 0.738 | 22.5 | >391.5 | 151.9 | 1.36±0.01 | 3.17±0.02 | 8.60±0.04 | 10.8±0.07 | 68.30±0.50 | 3-53 | K | 1-33 or 1D-33 |
| 222 | 0.019 ± 0.000 | 0.014 ± 0.002 | 0.017 ± 0.005 | 0.008 ± 0.003 | 0.7 | 0.9 | 0.4 | 1.36±0.08 | 3.96±0.11 | 5.16±0.12 | 2.25±0.04 | 1.92±0.01 | 3-53 | K | 3-20 |
| 253 | 0.055 ± 0.008 | 0.126 ± 0.018 | 0.109 ± 0.055 | 0.137 ± 0.005 | 2.3 | 2.0 | 2.5 | 1.15±0.03 | 5.99±0.11 | 6.11±0.11 | 2.66±0.03 | 4.25±0.11 | 1-58 | K | 3-20 |
| 269 | 0.030 ± 0.000 | >10 | >10 | >10 | >337.5 | >337.5 | >337.5 | 0.76±0.02 | 6.69±0.04 | 8.88±0.04 | 1.29±0.03 | Knocked out | 3-53 | K | 1-9 |
| 278 | 0.014 ± 0.007 | 0.307 ± 0.149 | 0.160 ± 0.018 | 0.245 ± 0.042 | 22.5 | 11.7 | 17.9 | 4.16±0.03 | 5.21±0.03 | 6.37±0.03 | 6.95±0.27 | 16.20±0.17 | 1-18 | K | 1-39 or 1D-39 |
| 281 | 0.005 ± 0.001 | 0.012 ± 0.000 | >10 | >10 | 2.5 | >2026.3 | >2026.3 | 0.97±0.03 | 0.68±0.02 | 4.42±0.03 | 3.54±0.08 | Knocked out | 3-7 | K | 2-24 |
| 316 | 0.018 ± 0.007 | 0.024 ± 0.005 | >10 | >10 | 1.4 | >563.6 | >563.6 | 4.81±0.05 | 4.96±0.06 | 5.17±0.08 | Knocked out | Knocked out | 1-2 | λ | 2-8 |
| 318 | 0.029 ± 0.008 | 0.185 ± 0.037 | 0.019 ± 0.008 | 0.083 ± 0.032 | 6.3 | 0.7 | 2.8 | 4.43±0.04 | 4.96±0.04 | 5.35±0.04 | 6.05±0.02 | 9.30±0.03 | 1-58 | K | 3-20 |
| 384 | 0.004 ± 0.001 | 0.005 ± 0.002 | >10 | >10 | 1.1 | >2398.4 | >2398.4 | 1.19±0.02 | 1.30±0.03 | 1.80±0.03 | 1.75±0.04 | Knocked out | 3-11 | K | 1-27 |
| 398 | 0.091 ± 0.004 | 0.180 ± 0.001 | >10 | >10 | 2.0 | >110.2 | >110.2 | 4.63±0.04 | 10.40±0.10 | 13.60±0.24 | Knocked out | Knocked out | 3-66 | λ | 2-8 |
| AZD1061 | 0.013 ± 0.003 | 0.012 ± 0.002 | 0.014 ± 0.002 | 0.007 ± 0.002 | 0.9 | 1.1 | 0.5 | 5.13±0.07 | 8.00±0.12 | 6.14±0.08 | 7.66±0.05 | 2.27±0.03 | N/A | N/A | N/A |
| AZD8895 | 0.005 ± 0.001 | 0.011 ± 0.002 | 0.046 ± 0.031 | 0.046 ± 0.016 | 2.2 | 8.9 | 8.8 | 2.18±0.04 | 2.96±0.05 | 3.56±0.06 | 8.50±0.05 | 7.35±0.15 | N/A | N/A | N/A |
| AZD7442 | 0.009 ± 0.000 | 0.007 ± 0.001 | 0.012 ± 0.001 | 0.006 ± 0.003 | 0.8 | 1.4 | 0.7 | N/A | N/A | N/A | N/A | N/A | N/A | N/A | N/A |
| REGN10987 | 0.032 ± 0.007 | 0.028 ± 0.003 | 0.007 ± 0.001 | 0.013 ± 0.002 | 0.9 | 0.2 | 0.4 | 1.35±0.03 | 1.61±0.03 | 1.84±0.04 | 1.39±0.05 | 1.38±0.02 | N/A | N/A | N/A |
| REGN10933 | 0.004 ± 0.002 | 0.014 ± 0.002 | 3.284 ± 2.014 | 6.177 ± 1.914 | 3.3 | 773.7 | 1455.2 | 0.97±0.02 | 1.99±0.02 | 1.70±0.01 | 1.92±0.02 | 308.7±10.0 | N/A | N/A | N/A |
| ADG10 | 0.006 ± 0.000 | 0.010 ± 0.001 | 0.011 ± 0.001 | 0.003 ± 0.000 | 1.8 | 1.9 | 0.5 | N/A | N/A | N/A | N/A | N/A | N/A | N/A | N/A |
| ADG20 | 0.004 ± 0.001 | 0.006 ± 0.000 | 0.010 ± 0.001 | 0.009 ± 0.000 | 1.4 | 2.5 | 2.2 | N/A | N/A | N/A | N/A | N/A | N/A | N/A | N/A |
| ADG30 | 0.007 ± 0.002 | 0.016 ± 0.001 | 0.029 ± 0.003 | 0.002 ± 0.001 | 2.5 | 4.4 | 0.3 | N/A | N/A | N/A | N/A | N/A | N/A | N/A | N/A |
| LY-CoV555 | 0.006 ± 0.002 | 0.009 ± 0.000 | >10 | >10 | 1.5 | >1545.3 | >1545.3 | N/A | N/A | N/A | N/A | N/A | N/A | N/A | N/A |
| LY-CoV16 | 0.034 ± 0.007 | 3.225 ± 1.030 | >10 | >10 | 1.5 | >291.2 | >291.2 | N/A | N/A | N/A | N/A | N/A | N/A | N/A | N/A |
| S309 | 0.040 ± 0.005 | 0.078 ± 0.069 | 0.082 ± 0.002 | 0.076 ± 0.014 | 1.5 | 2.0 | 1.9 | N/A | N/A | N/A | N/A | N/A | N/A | N/A | N/A |
| 222H+222L | 0.017 ± 0.001 | 0.011 ± 0.002 | 0.016 ± 0.001 | 0.009 ± 0.000 | 0.6 | 1.0 | 0.5 | N/A | N/A | N/A | N/A | N/A | 3-53 | K | 3-20 |
| 150H+222L | 0.016 ± 0.003 | 0.010 ± 0.001 | 0.007 ± 0.001 | 0.003 ± 0.000 | 0.6 | 0.4 | 0.2 | N/A | N/A | N/A | N/A | N/A | 3-53 | K | 3-20 |
| 158H+222L | 0.033 ± 0.003 | 0.014 ± 0.001 | 0.056 ± 0.015 | 0.019 ± 0.000 | 0.4 | 1.7 | 0.6 | N/A | N/A | N/A | N/A | N/A | 3-53 | K | 3-20 |
| 175H+222L | >10 | 0.399 ± 0.012 | >10 | >10 | <0.04 | N/A | N/A | N/A | N/A | N/A | N/A | N/A | 3-53 | K | 3-20 |
| 269H+222L | 0.552 ± 0.085 | 0.024 ± 0.001 | >10 | >10 | 0.04 | >18.1 | >18.1 | N/A | N/A | N/A | N/A | N/A | 3-53 | K | 3-20 |

### TableS3A

| Convalescent plasma | FRNT50 (Recprocal plasma dilution) | | Victoria/P.1 ratio |
|---|---|---|---|
| | Victoria | P.1 | |
| Convalescent 1 | 61 | 25 | 2.5 |
| Convalescent 2 | 689 | 49 | 14.2 |
| Convalescent 3 | 526 | 173 | 3.0 |
| Convalescent 4 | 409 | 356 | 1.1 |
| Convalescent 5 | 369 | 105 | 3.5 |
| Convalescent 6 | 1270 | 556 | 2.3 |
| Convalescent 7 | 274 | 80 | 3.4 |
| Convalescent 8 | 633 | 223 | 2.8 |
| Convalescent 9 | 667 | 126 | 5.3 |
| Convalescent 10 | 124 | 34 | 3.7 |
| Convalescent 11 | 102 | 30 | 3.4 |
| Convalescent 12 | 339 | 74 | 4.6 |
| Convalescent 13 | 331 | 240 | 1.4 |
| Convalescent 14 | 438 | 95 | 4.6 |
| Convalescent 15 | 6397 | 3261 | 2.0 |
| Convalescent 16 | 44 | 39 | 1.1 |
| Convalescent 17 | 1115 | 87 | 12.8 |
| Convalescent 18 | 242 | 64 | 3.8 |
| Convalescent 19 | 29 | 20 | 1.4 |
| Convalescent 20 | 154 | 136 | 1.1 |
| Convalescent 21 | 487 | 165 | 3.0 |
| Convalescent 22 | 438 | 241 | 1.8 |
| Convalescent 23 | 381 | 83 | 4.6 |
| Convalescent 24 | 1647 | 390 | 4.2 |
| Convalescent 25 | 913 | 322 | 2.8 |
| Convalescent 26 | 1880 | 825 | 2.3 |
| Convalescent 27 | 1464 | 206 | 7.1 |
| Convalescent 28 | 361 | 81 | 4.4 |
| Convalescent 29 | 2859 | 1010 | 2.8 |
| Convalescent 30 | 1109 | 477 | 2.3 |
| Convalescent 31 | 811 | 274 | 3.0 |
| Convalescent 32 | 395 | 130 | 3.0 |
| Convalescent 33 | 1144 | 207 | 5.5 |
| Convalescent 34 | 676 | 201 | 3.4 |

### TableS3B

| B.1.1.7 plasma | Day post symptom onset | FRNT50 (Recprocal plasma dilution) | | Victoria/P.1 ratio |
|---|---|---|---|---|
| | | Victoria | P.1 | |
| B.1.1.7 P3 | 11 | 440 | 116 | 3.8 |
| B.1.1.7 P4 | 18 | 136884 | 48440 | 2.8 |
| B.1.1.7 P5 | 45 | 1506 | 787 | 1.9 |
| B.1.1.7 P6 | 31 | 370 | 259 | 1.4 |
| B.1.1.7 P7 | 33 | 2250 | 267 | 8.4 |
| B.1.1.7 P8 | 25 | 2999 | 2261 | 1.3 |
| B.1.1.7 P9 | 26 | 970 | 861 | 1.1 |
| B.1.1.7 P10 | 18 | 3735 | 861 | 4.3 |
| B.1.1.7 P11 | 24 | 2193 | 1116 | 2.0 |
| B.1.1.7 P13 | 29 | <20 | 67 | <0.3 |
| B.1.1.7 P14 | 21 | 1700 | 168 | 10.1 |
| B.1.1.7 P15 | 16 | 168 | 414 | 0.4 |

### Table S4

| Vaccine samples | Day Post-boost | FRNT50 (Recprocal serum dilution) | | Victoria/P.1 ratio |
|---|---|---|---|---|
| | | Victoria | P.1 | |
| Pfizer1 | 7 | 1149 | 396 | 2.9 |
| Pfizer2 | 7 | <20 | 36 | <0.6 |
| Pfizer3 | 7 | 1727 | 698 | 2.5 |
| Pfizer4 | 8 | 2234 | 712 | 3.1 |
| Pfizer5 | 7 | 3016 | 1033 | 2.9 |
| Pfizer6 | 7 | 1521 | 302 | 5.0 |
| Pfizer7 | 7 | 609 | 294 | 2.1 |
| Pfizer8 | 7 | 4340 | 2119 | 2.0 |
| Pfizer9 | 7 | 1467 | 361 | 4.1 |
| Pfizer10 | 7 | 1757 | 343 | 5.1 |
| Pfizer11 | 7 | 860 | 424 | 2.0 |
| Pfizer12 | 7 | 1749 | 452 | 3.9 |
| Pfizer13 | 7 | 1851 | 669 | 2.8 |
| Pfizer14 | 7 | 407 | 294 | 1.4 |
| Pfizer15 | 8 | 1285 | 571 | 2.3 |
| Pfizer16 | 8 | 1286 | 311 | 4.1 |
| Pfizer17 | 8 | 1810 | 304 | 6.0 |
| Pfizer18 | 8 | 1198 | 282 | 4.3 |
| Pfizer19 | 8 | 466 | 229 | 2.0 |
| Pfizer20 | 8 | 1539 | 693 | 2.2 |
| Pfizer21 | 9 | 184 | 52 | 3.5 |
| Pfizer22 | 11 | 1061 | 491 | 2.2 |
| Pfizer23 | 12 | 1658 | 355 | 4.7 |
| Pfizer24 | 12 | 1155 | 569 | 2.0 |
| Pfizer25 | 15 | 8092 | 5029 | 1.6 |
| AstraZeneca 1 | 28 | 495 | 265 | 1.9 |
| AstraZeneca 2 | 28 | 580 | 429 | 1.4 |
| AstraZeneca 3 | 28 | 253 | <20 | >12.6 |
| AstraZeneca 4 | 28 | 183 | 102 | 1.8 |
| AstraZeneca 5 | 28 | 432 | 215 | 2.0 |
| AstraZeneca 6 | 28 | 764 | 111 | 6.9 |
| AstraZeneca 7 | 28 | 133 | 29 | 4.5 |
| AstraZeneca 8 | 28 | 257 | 116 | 2.2 |
| AstraZeneca 9 | 28 | 501 | 97 | 5.2 |
| AstraZeneca 10 | 28 | 357 | 133 | 2.7 |
| AstraZeneca 11 | 14 | 334 | 115 | 2.9 |
| AstraZeneca 12 | 14 | 250 | 94 | 2.7 |
| AstraZeneca 13 | 14 | 122 | 40 | 3.1 |
| AstraZeneca 14 | 14 | 212 | 110 | 1.9 |
| AstraZeneca 15 | 14 | 789 | 281 | 2.8 |
| AstraZeneca 16 | 14 | 538 | 181 | 3.0 |
| AstraZeneca 17 | 14 | 1159 | 359 | 3.2 |
| AstraZeneca 18 | 14 | 353 | 85 | 4.1 |
| AstraZeneca 19 | 14 | 975 | 382 | 2.6 |
| AstraZeneca 20 | 14 | 169 | 74 | 2.3 |
| AstraZeneca 21 | 14 | 155 | 87 | 1.8 |
| AstraZeneca 22 | 14 | 152 | 98 | 1.5 |
| AstraZeneca 23 | 14 | 126 | 67 | 1.9 |
| AstraZeneca 24 | 14 | 293 | 151 | 1.9 |
| AstraZeneca 25 | 14 | 94 | 25 | 3.8 |

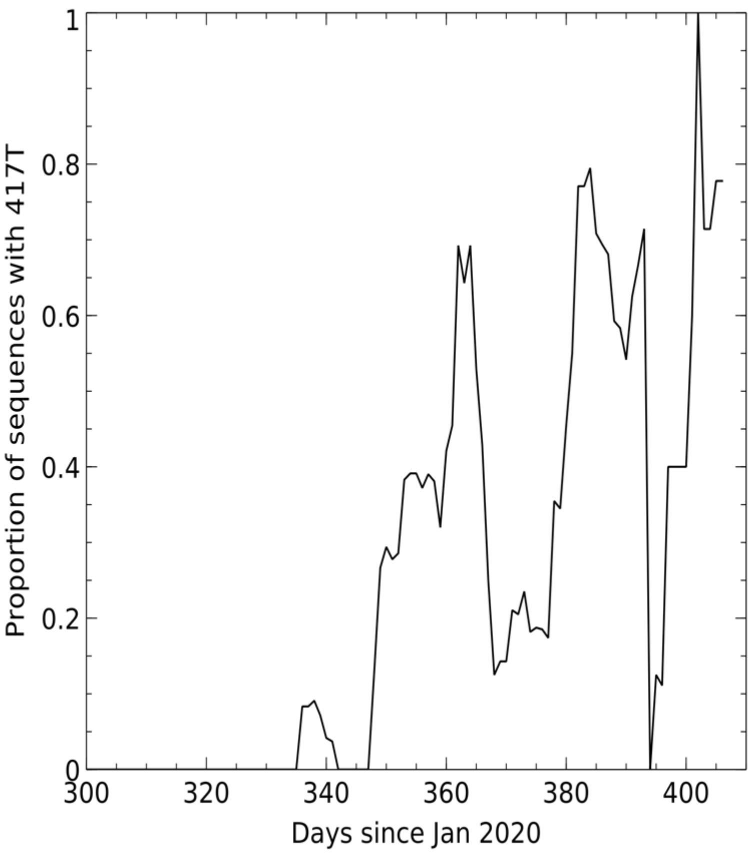


Sliding 7-day window depicting proportion of sequences containing K417T

Figure S1

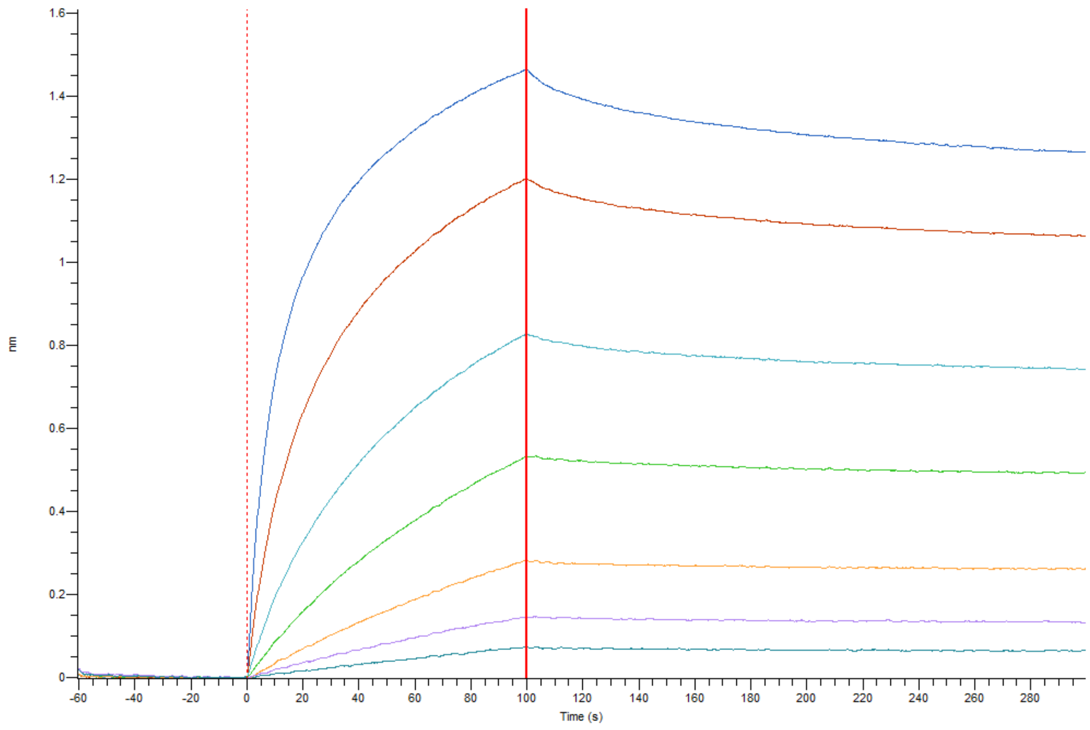


BLI titration for the attachment and dissociation of ACE2 from P.1 RBD attached to the tip

Fig. S2
